# FoXSight: Facilitating Computational Hit Identification in Crystallographic Fragment Screening Experiments

**DOI:** 10.64898/2026.11.01.763443

**Authors:** Thorben Schulze, Sven Falke, Sebastian Günther, Alke Meents, Matthias Rarey

## Abstract

X-ray crystallographic fragment screening reveals how small molecules bind to proteins, providing starting points for drug design. Identifying these weak binding events remains difficult when electron density signals are noisy, partially occupied or obscured by solvent, and campaigns may comprise thousands of datasets. We present FoXSight, a computational method for hit identification in crystallographic compound screening. FoXSight applies Difference-of-Gaussian (DoG) filtering to multi-reference subtraction maps to detect recurring, target-specific density features associated with bound ligands. Its output maps remain visually interpretable and support ligand modeling. Across six annotated campaigns comprising 2,715 datasets, the top 10% of each FoXSight ranking contained 134 of 264 confirmed hits. FoXSight served as the primary hit identification tool for a human Cathepsin L campaign with non-isomorphous crystals, where the output of established methods was impractical to inspect systematically. The FoXSight computations for its 1,481 datasets took 47 minutes on 75 CPU cores.

---

Structure-based fragment screening has become a cornerstone in early-stage drug discovery. It enables researchers to investigate how small molecules (fragments or compounds) interact with pharmaceutically relevant target proteins, providing crucial starting points for lead development [1–4]. Various experimental methods are used for fragment screening, including NMR spectroscopy and biosensor-based approaches, which are especially well suited to detect binding events. However, these methods often do not reveal the binding mode or structural orientation of the fragment. In contrast, X-ray crystallography can not only confirm binding but also provide precise, atomic detail of how a fragment interacts with the protein, which is essential for rational drug design. Recent advances in data processing and density analysis have improved the detection of low-occupancy binding events [5, 6].

The demand for new therapeutics in the face of global health threats—such as highly pathogenic RNA viruses (e.g., SARS-CoV-2) and the spread of antibiotic-resistant bacteria—has propelled high-throughput crystallographic screening to the forefront. Recent technological advances at synchrotron sources allow the collection of several hundreds of crystal datasets within a single day. Meanwhile, optimized fragment libraries—often encompassing more than one thousand compounds—are readily available for such large-scale screens [7, 8]. The SARS-CoV-2 main-protease screen of repurposing libraries illustrates the scale of such campaigns [3]. Recent advances in serial crystallography also extend high-throughput fragment screening to room temperature, as demonstrated for the fosfomycin-resistance protein FosAKP [9]. These capabilities generate an enormous amount of diffraction data, presenting significant computational challenges for accurate and efficient hit identification, particularly for only weakly binding fragments.

Hit identification in such large screening campaigns is non-trivial. Fragments typically have a low molecular weight (below 300 Da) and low affinity, often leading to weak and ambiguous electron density signals. Sensitive and specific computational tools that can analyze large datasets efficiently are thus essential. Among current tools, PanDDA [10] is widely used. It analyzes multiple datasets statistically to construct a “true background” electron density, helping to reveal otherwise hidden binding events and achieving a high detection rate. However, PanDDA heavily relies on a sufficient amount of isomorphous data to build the background. This requirement can introduce a skew toward hit states in the background or a non-stable background leading to numerous false negative or positive hits, respectively. Other computational tools can aid in hit identification, such as cluster4x [11], which groups similar datasets prior to hit finding with PanDDA, and Xtrapol8, which is based on pairwise dataset comparison to enhance weak differences to increase the visibility of the ligand density and mainly applied in time-resolved crystallography [12]. The growing number of such methods reflects the importance of X-ray crystallographic fragment screening and the necessity for more reliable and efficient computational hit identification approaches.

To address these limitations, we developed FoXSight (Fragment Observer for X-ray Screenings Incorporating Gaussian Highlighting Techniques), a computational tool for automated hit identification across crystallographic datasets ranging from small compound-validation experiments to large-scale fragment screening campaigns. FoXSight evaluates each target dataset individually against a target-specific ensemble of suitable background references and applies Difference-of-Gaussian (DoG) filtering to enhance weak, target-specific density features associated with bound ligands, including cases with non-isomorphism (Fig. 1). Rather than replacing PanDDA, FoXSight provides a complementary perspective on ligand-associated density and can thereby support hit identification and increase confidence in challenging or ambiguous cases.

**Fig. 1.**
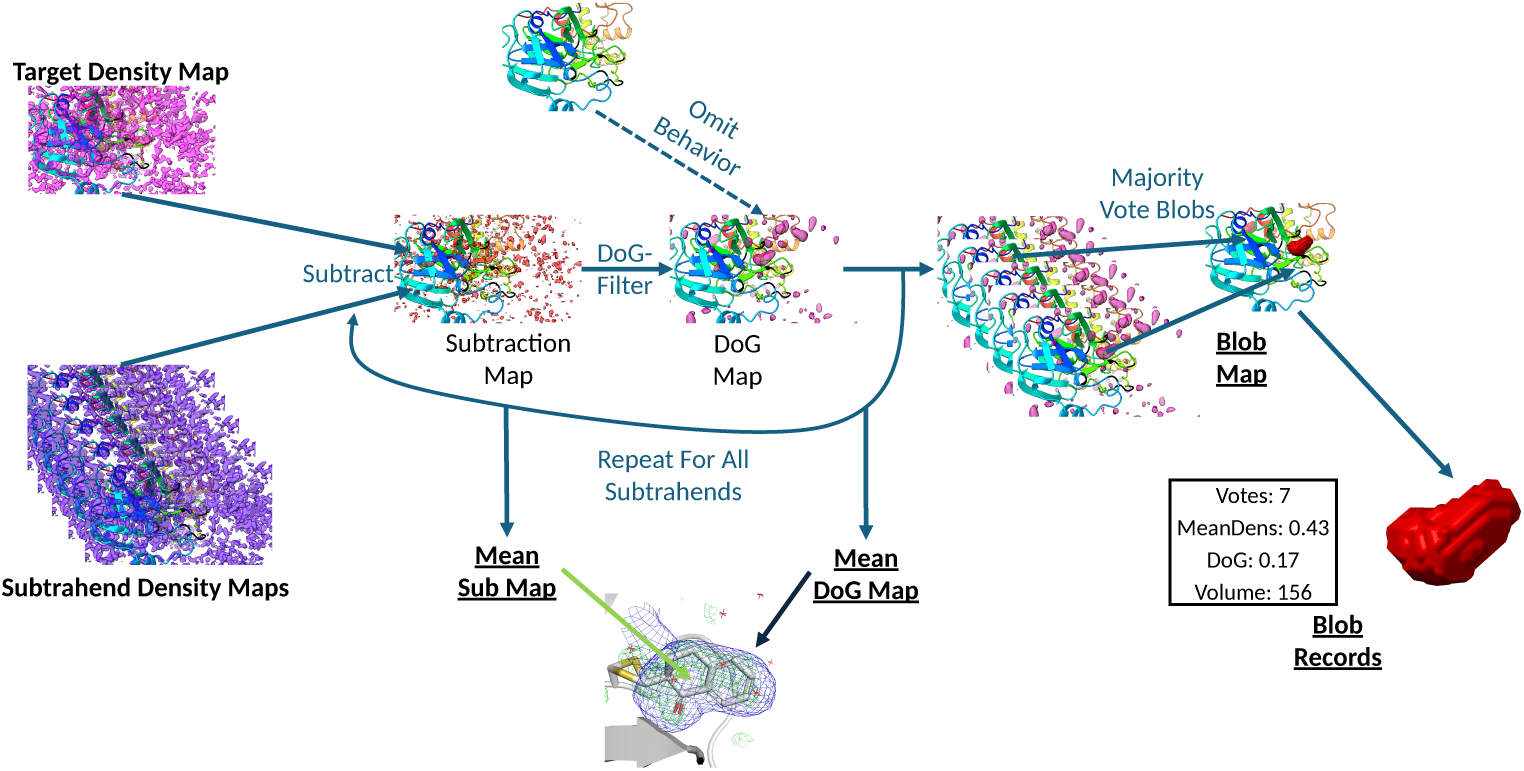
Overview of the FoXSight workflow. For each target dataset, the 2*mF*o − *DF*c density map is separately compared with selected subtrahend maps derived from reference datasets. Each target-subtrahend pair yields a subtraction map, with density surrounding the protein model and, optionally, solvent molecules omitted before DoG filtering. Difference-of-Gaussian (DoG) filtering suppresses diffuse map variation and enhances localized residual electron-density features. Contiguous regions close to local maxima are segmented into blobs using a breadth-first search. Blobs are compared across subtraction maps, and similar blobs from other maps contribute consensus votes. The originating map does not vote for its own blobs. The resulting blob map highlights spatially recurring features, while the associated records summarize votes, mean density, DoG response and blob volume. The meanDoG map represents the broader spatial extent of localized features, whereas the mean subtraction map localizes the strongest residual density differences more directly.

## 1 Results

### 1.1 Datasets

FoXSight was evaluated on screening datasets for seven protein targets: the SARS-CoV-2 *main protease (M^pro^)* [3], the *nuclear auto-antigen SP-100 (SP100)* [13], the *bromodomain BRD1* [14], the *demethylase JMJD2D* [15], the *bromodomain BAZ2B* [16], *human JMJD1B* [17], and *human Cathepsin L (CatL)*. The five public fragment-screening datasets provide PanDDA-derived hit annotations for the enrichment analysis. The M^pro^ campaign is considered separately for recovery of reported hits and comparison of representative maps, because its original hit identification did not use a uniform PanDDA analysis across the campaign. FoXSight served as the primary hit identification method for the Cathepsin L campaign.

### 1.2 Validation

To validate FoXSight, we examined the recovery of reported hits from previous screening campaigns, including examples from the original PanDDA publication. We also used FoXSight as the primary hit identification method for a large screening campaign for human Cathepsin L. The European Fragment Screening Library (EFSL) [18], comprising 1,056 fragments, was screened in this campaign. The crystals belonged to space group P1, contained four protein chains in the asymmetric unit, and showed variation in unit-cell parameters and protein conformation (Supplementary Figures S1 and S2). Unless stated otherwise, all experiments used the same numerical parameters, with both bounding-box options enabled (Supplementary Table S3). For SP100, BRD1, BAZ2B, JMJD2D and JMJD1B, 100 randomly selected datasets formed the candidate reference pool. For the other campaigns, all known apo structures were considered as candidate references. The enrichment analysis comprised 2,715 datasets, including all 226 JMJD2D datasets. The runtime benchmark additionally included 25 M^pro^ datasets, for a total of 2,740 analyses. For visualization, we used PyMOL [19] and a consistent carve radius of 2.5 Å. Unless stated otherwise, conventional 2*mF*_o_ *− DF*_c_ maps were displayed at 1.0*σ* and *mF*_o_ *DF*_c_ maps at 3.0*σ*. PanDDA event maps were displayed at 2.0*σ* and Z-maps at raw Z-scores of 4, the level used in the original PanDDA publication [10]. FoXSight meanDoG maps were displayed at 3.0*σ* and mean subtraction maps at 2.0*σ*. These contour levels were kept consistent across the displayed comparisons and are distinct from detection thresholds. Because carving is applied around the modeled atoms, it also limits the displayed extent of spatially broad features, particularly of the meanDoG maps.

### 1.3 Blob-Based Scoring Scheme for Presorting Hit Candidates

We introduced an empirical score to rank datasets for manual inspection while retaining the maps for assessment of each candidate. It combines consensus votes, mean DoG response, volume and mean density, down-weights blobs much larger than a typical fragment, and normalizes their sum by the overall target–reference difference (Methods). The enrichment analysis covers five public campaigns with PanDDA-derived annotations and the Cathepsin L campaign, whose labels were established in a FoXSight-led workflow (Table 1 and Fig. 2). M^pro^ was excluded because a uniform campaign-wide PanDDA reference analysis was unavailable. Normalization and volume weighting increased the confirmed hits in the highest-ranked 10% from 105 to 134 of 264 compared with the blob sum alone (Supplementary Table S4). Gains occurred mainly in Cathepsin L (54 to 74), BRD1 (21 to 26) and JMJD1B (12 to 17), while JMJD2D decreased from 9 to 7. Parameter-sensitivity and campaign-held-out analyses are reported in the Supplementary Information.

**Fig. 2.**
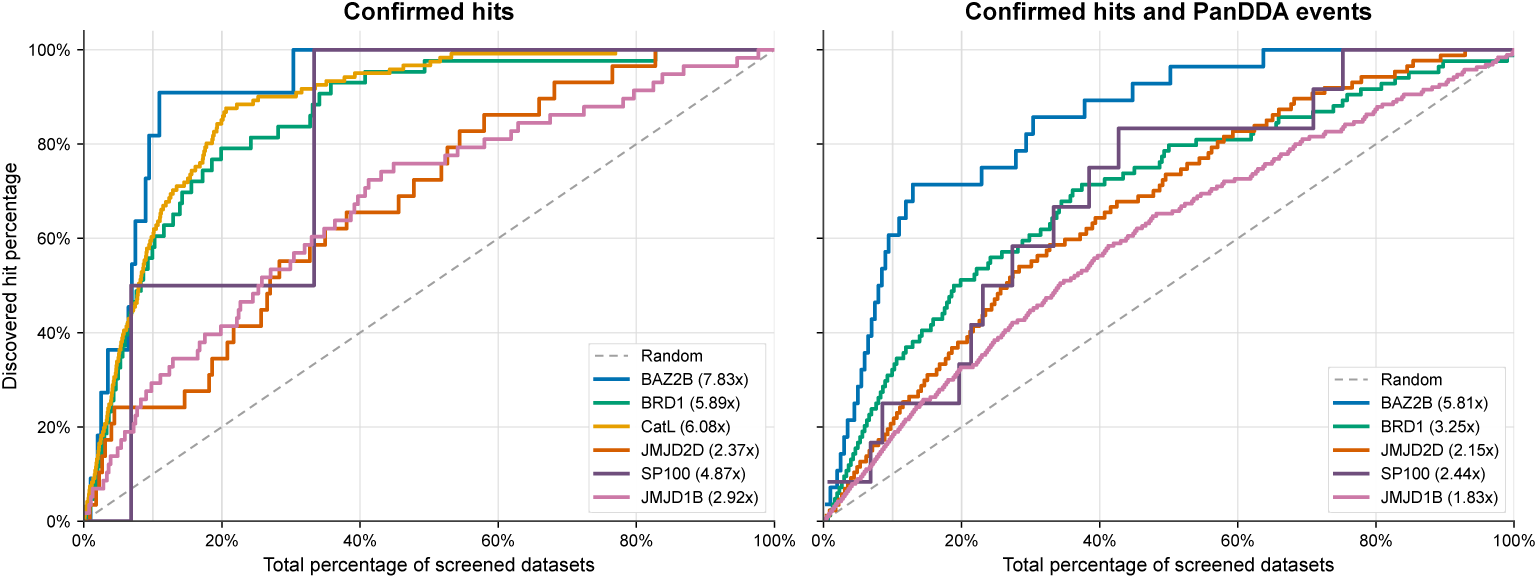
Enrichment of the empirical FoXSight score for prioritizing datasets. Blob contributions combine squared votes, mean DoG response, mean target density and a volume weight that decreases above 300 Å^3^. Their sum is divided by the 1.5th power of the subtraction residual. The left panel evaluates confirmed-hit annotations. The right panel additionally treats every annotated PanDDA event as a positive, providing a broader comparison with event annotations rather than an independently validated ligand-hit set. Some events can represent genuine ligand density not reported as a confirmed hit, including binding at crystal contacts. Cathepsin L has no event annotations and appears only in the left panel. The larger positive fraction in the right panel lowers the maximum attainable EF10, to approximately 2.0 for JMJD1B. Legend values are EF10.

**Table 1.** Confirmed-hit enrichment of the FoXSight dataset score. The score is blobscore*_d_/R*^1.5^, with the volume weight of Eq. (5). Datasets were ranked by decreasing score. Top-10% and top-50% recovery counts confirmed-hit datasets among the first ⌈*f N* ⌉ ranks relative to the total confirmed-hit count. EF10 is the hit-rate enrichment factor at 10%. Recovery AUC is the area under the cumulative recovery curve against the inspected dataset fraction. It is not ROC AUC. AP is average precision and should be interpreted alongside the campaign’s hit prevalence. Labels derive from the available campaign annotations. Cathepsin L labels were established in a FoXSight-led workflow. Results for the unnormalized, unweighted blob sum are in Supplementary Table S4.

| Target | *N* | Full hits | Top 10% recovery (*n*/*N*) | Top 50% recovery (*n*/*N*) | EF10 | Recovery AUC | AP |
|---|---|---|---|---|---|---|---|
| BAZ2B | 201 | 11 | 9/11 | 11/11 | 7.83 | 0.921 | 0.455 |
| BRD1 | 302 | 43 | 26/43 | 42/43 | 5.89 | 0.859 | 0.722 |
| JMJD2D | 226 | 29 | 7/29 | 21/29 | 2.37 | 0.682 | 0.310 |
| SP100 | 117 | 2 | 1/2 | 2/2 | 4.87 | 0.803 | 0.088 |
| JMJD1B | 388 | 58 | 17/58 | 44/58 | 2.92 | 0.678 | 0.344 |
| CatL | 1,481 | 121 | 74/121 | 117/121 | 6.08 | 0.881 | 0.481 |
| Pooled recovery | 2,715 | 264 | 134/264 | 237/264 | – | – | – |

### 1.4 Cathepsin L

The fragment-screening campaign for human Cathepsin L was carried out at the MX beamline P11 of PETRA III at DESY using the 1,056-fragment EFSL [18]. Diffraction data were collected from 2,260 single crystals at cryogenic temperatures. Of these, 1,481 datasets, including 42 apo datasets, passed the complete DIMPLE pipeline [20] and were potentially suitable for FoXSight analysis and model building. Purification and crystallization conditions have been reported previously [21]. Data collection and refinement statistics for the four representative hits shown in Fig. 3 are provided in Supplementary Table S2.

**Fig. 3.**
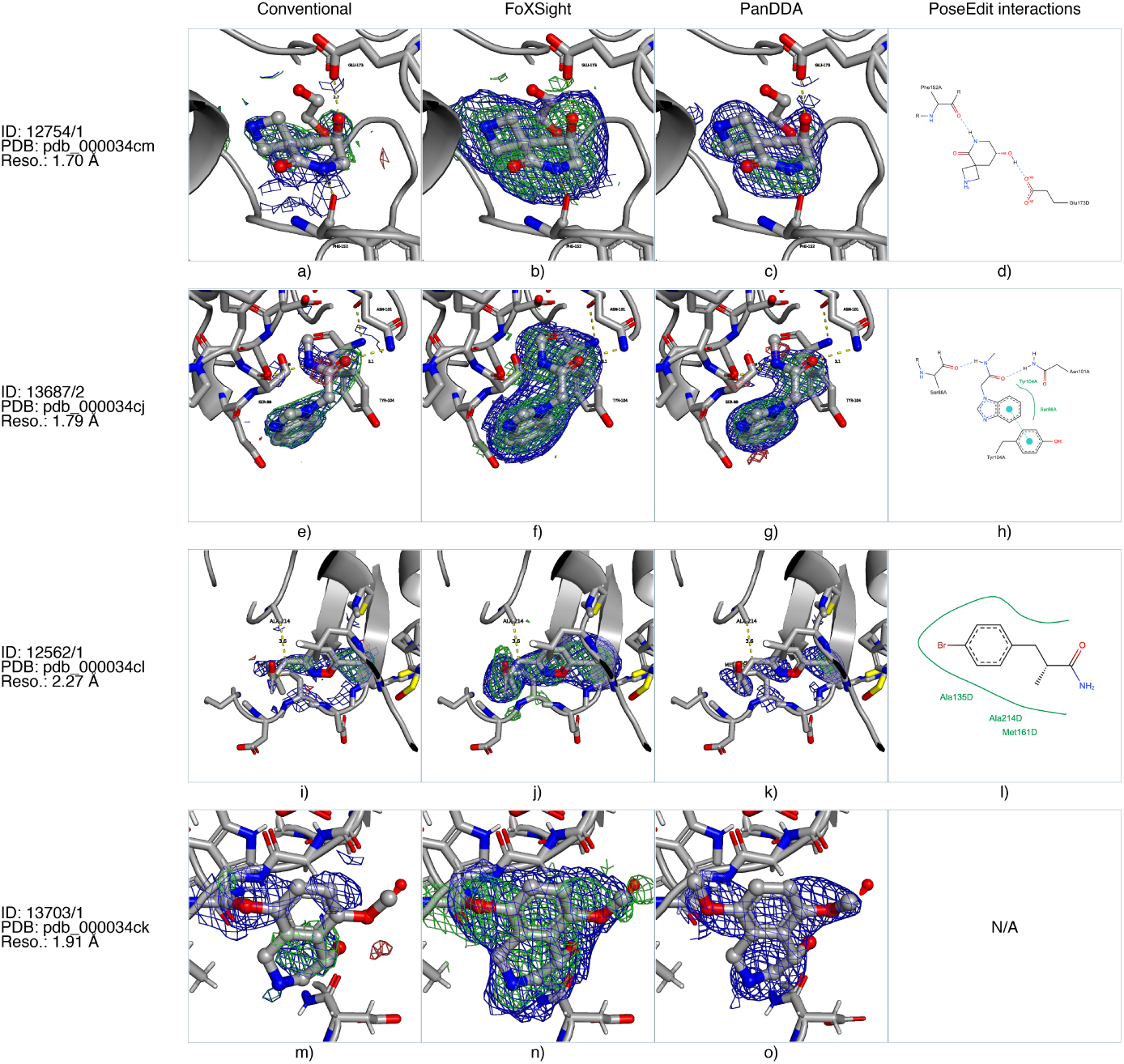
Representative human Cathepsin L hit maps and protein–ligand interaction diagrams. Rows show datasets 12754/1 at 1.70 Å (a–d), 13687/2 at 1.79 Å (e–h), 12562/1 at 2.27 Å (i–l), and 13703/1 at 1.91 Å (m–o). Full crystal, compound, and PDB/deposition identifiers are listed in Supplementary Table S1. Conventional 2*mF*o − *DF*c maps are blue and *mF*o − *DF*c maps green/red. FoXSight meanDoG maps are blue and mean subtraction maps green. PanDDA event maps are blue and Z-maps green/red. FoXSight and PanDDA reveal more coherent ligand-associated density than conventional maps in these examples. In panel n, the ligand is shown with the alternative orientation of one ether moiety suggested by the FoXSight maps. PoseEdit [22] diagrams summarize interactions for the refined poses. For 13703/1, no interactions met the PoseEdit distance and angle thresholds.

These 1,481 datasets formed the ranked FoXSight cohort, with the 42 apo structures as candidate references. Its 121 confirmed-hit datasets correspond to 121 distinct compound database identifiers in the campaign hit list. FoXSight maps and blob-based scoring guided hit identification, model building and refinement. For several hits, conventional 2*mF*_o_ *DF*_c_ and *mF*_o_ *DF*_c_ maps were ambiguous, whereas the meanDoG map indicated the broader ligand-associated region and the mean subtraction map localized the strongest residual features.

The CatL data were also processed with PanDDA after cluster4x subdivision to address crystallographic heterogeneity (Supplementary Figures S1 and S2). In a qualitative assessment of practical manual inspectability, the resulting clusters were graded as good, okay, medium, bad or really bad. The good, okay and medium categories contained 46 (38%) of the 121 known hit datasets. This assessment describes the practical triage achieved in that workflow, rather than the sensitivity of an exhaustive PanDDA analysis (Supplementary Results).

FoXSight and the available PanDDA maps generally supported similar ligand placements. For dataset 13703/1, the FoXSight maps suggested a different orientation of one ether moiety that remained chemically plausible and compatible with the displayed density (Fig. 3). Such differences can guide evaluation of alternative poses, but greater map coherence alone does not establish a more accurate atomic model.

### 1.5 Case Study: BAZ2B

The BAZ2B dataset [16] provides a well-characterized public dataset for evaluating FoXSight against conventional electron density maps and PanDDA. It contains several challenging binding events in which ligand density is weak, partially obscured by solvent, or difficult to distinguish from the ground state [10], allowing the complementary information provided by the FoXSight maps to be examined directly. Representative examples are shown in Fig. 4, and further examples in Supplementary Figure S3.

**Fig. 4.**
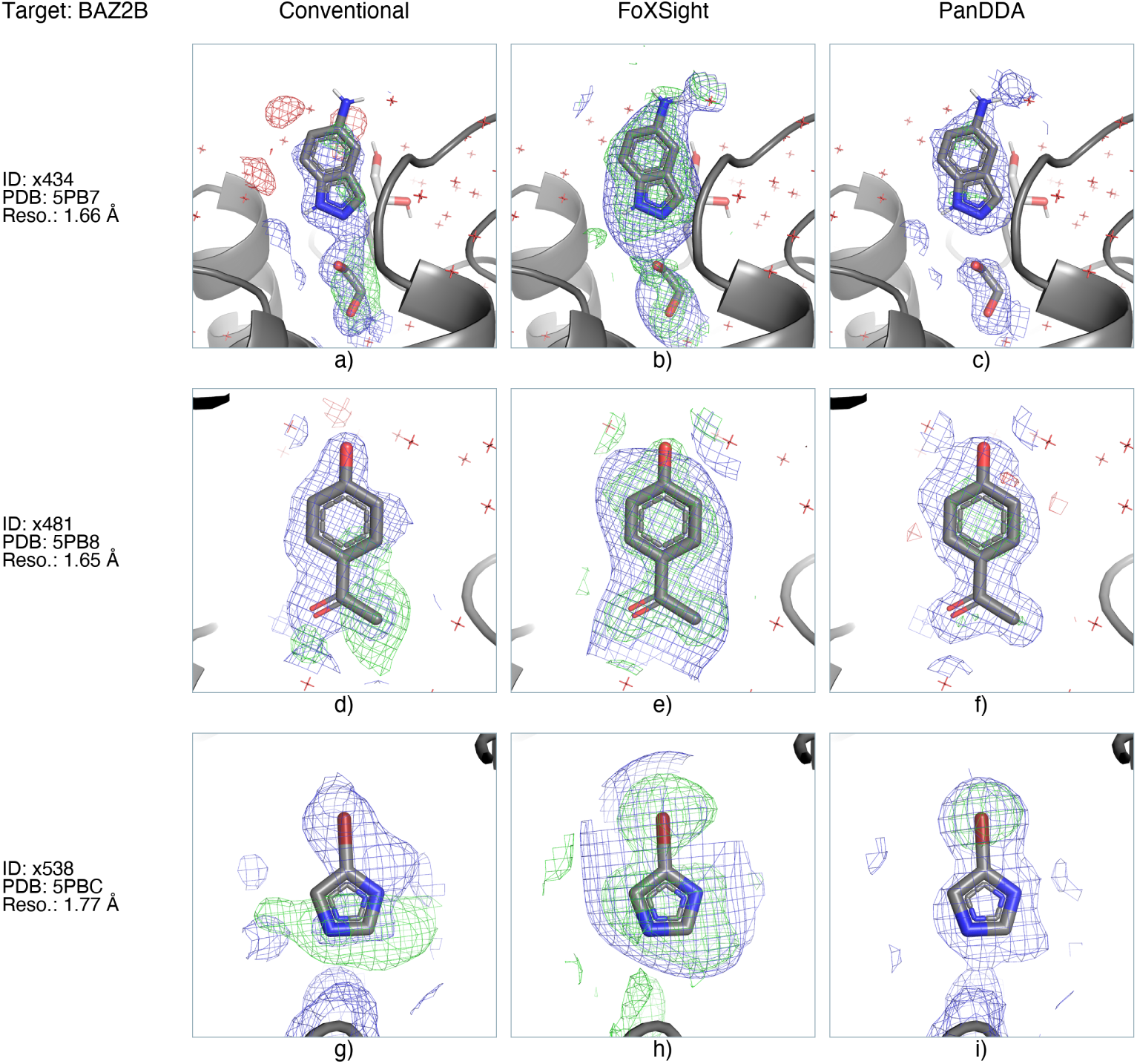
Representative BAZ2B map comparisons. Columns show conventional 2*mF*o − *DF*c (blue) and *mF*o − *DF*c maps (green/red), FoXSight meanDoG (blue) and mean subtraction maps (green), and PanDDA event (blue) and Z-maps (green/red). For x434, the conventional maps (a) show fragmented density around the ligand, the FoXSight maps (b) cover the ligand together with an adjacent modeled solvent molecule, and the PanDDA event map (c) outlines the ligand. For x481 and x538, the meanDoG map outlines a broader region and the mean subtraction map localizes the strongest differences near the modeled ligand atoms (panels e and h). Contours are specified in the Validation subsection.

Dataset x538 provides a particularly illustrative low-occupancy example. In the original PanDDA analysis, the bromine-containing fragment was superposed with a partially occupied ethylene glycol ground state. Therefore, the conventional electron density maps were difficult to interpret [10]. The PanDDA event map revealed the ligand-associated density, while anomalous difference density at the bromine atom provided independent experimental support for the ligand assignment. FoXSight similarly highlights the binding event: the meanDoG map outlines the broader ligand-associated region, whereas the mean subtraction map localizes the strongest density differences close to the ligand atoms. The independently observed bromine signal demonstrates that FoXSight can recover weak ligand-associated features despite competing solvent density.

### 1.6 Case Study: M^pro^

The crystallographic repurposing screen for M^pro^ revealed chemically diverse ligands with different molecular dimensions and binding modes [3]. The present analysis included a subset of 25 hit datasets reported in that study. FoXSight recovered ligand-associated density for all 25 hits. We selected representative examples to examine whether FoXSight detects spatially extended ligand-associated density when conventional or available PanDDA maps provide incomplete or fragmented coverage at the applied contour levels (Fig. 5 and Supplementary Figure S5). These comparisons assess individual binding events rather than campaign-wide enrichment, because the original campaign used heterogeneous hit-identification procedures without a uniform PanDDA analysis.

**Fig. 5.**
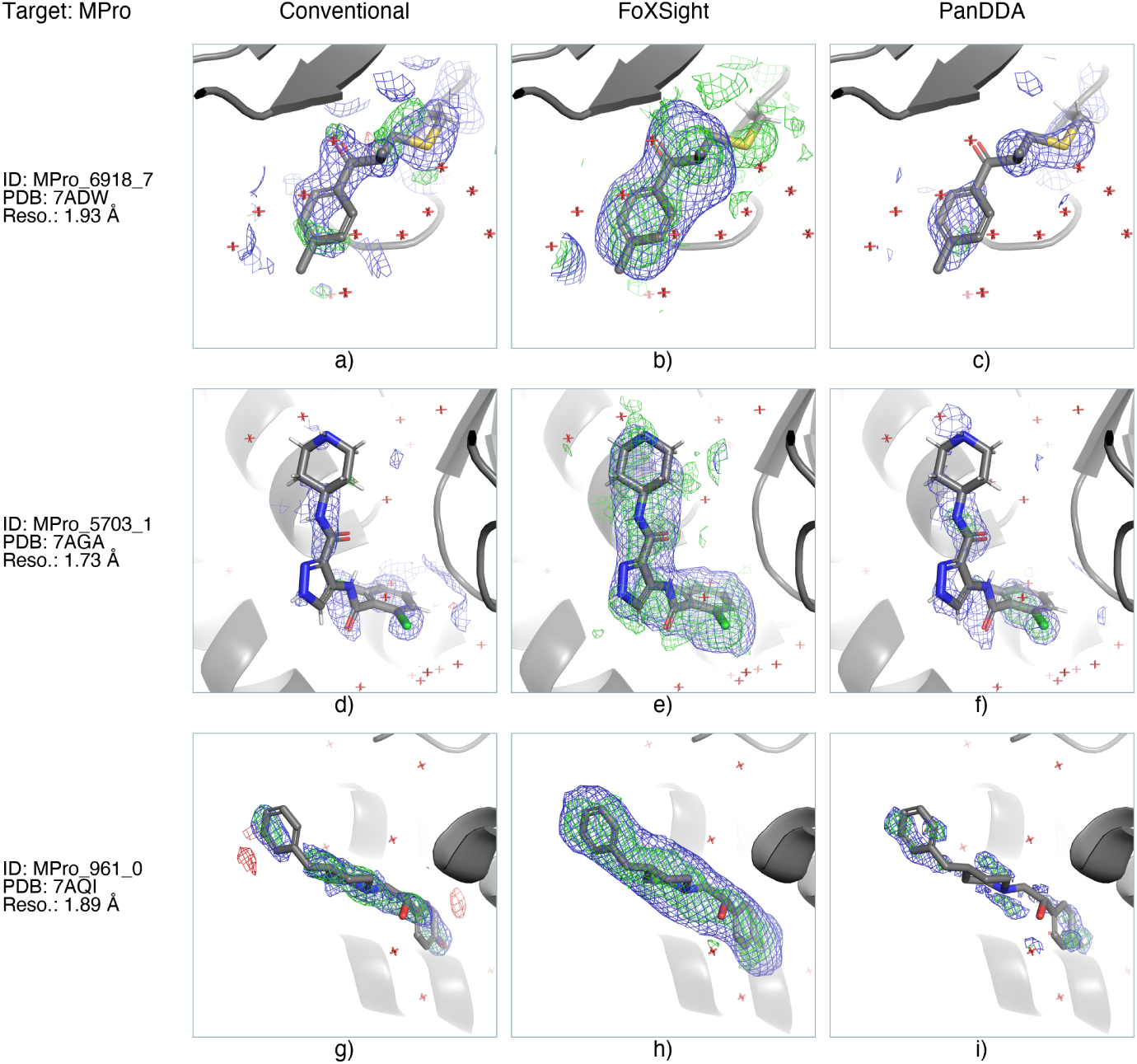
Representative SARS-CoV-2 M^pro^ hit maps. Columns show conventional 2*mF*o − *DF*c (blue) and *mF*o − *DF*c (green/red) maps, FoXSight meanDoG (blue) and mean subtraction (green) maps, and available PanDDA event (blue) and Z-maps (green/red). Rows correspond to dataset MPro 6918 7 (PDB 7ADW: a–c), MPro 5703 1 (PDB 7AGA: d–f), and MPro 961 0 (PDB 7AQI: g–i). At the stated contour levels, the FoXSight meanDoG maps form coherent envelopes around substantial portions of these ligands, while the mean subtraction maps localize the strongest residual density nearer the modeled atoms. The conventional and PanDDA maps show more fragmented coverage in some of these examples.

Several examples suggest that FoXSight is particularly helpful for spatially demanding compounds. For the datasets 7ADW, 7AGA and 7AQI, the meanDoG maps form coherent density envelopes that cover large portions of the ligand structures, whereas the corresponding conventional and, in some cases, PanDDA maps appear more fragmented at the chosen contour level. The mean subtraction maps complement this broader signal by localizing the strongest density differences closer to the ligand atoms. Although these selected examples do not prove a general advantage of FoXSight over other approaches for larger compounds, they illustrate that FoXSight can facilitate visual interpretation when ligand-associated density is distributed across a spatially demanding molecular structure.

### 1.7 Reproduction of cluster4x Hits

In the cluster4x study [11], grouping datasets according to crystallographic similarity before PanDDA analysis revealed several hits that had not been identified in the original PanDDA analysis. We examined whether FoXSight could recover the two additional BAZ2B hits (x447 and x557) without cluster-informed reference selection. Despite using randomly selected reference datasets, FoXSight produced clearly interpretable density features for both hits in the corresponding binding sites (Supplementary Figure S4). These hits, like the additional cluster4x hits in JMJD2D, are included in the enrichment analysis (Table 1).

## 2 Discussion

Across the analyzed publicly available and in-house crystallographic screening datasets, FoXSight consistently detected and localized electron-density features associated with ligand binding. The method combines multi-reference subtraction, DoG filtering, blob detection, and consensus voting. These features allow FoXSight to identify meaningful electron-density features related to ligand binding. Therefore, FoXSight can be used to convert a large collection of crystallographic datasets into a ranked and visually interpretable set of binding events, which enables efficient identification of the most likely binders. Importantly, FoXSight’s prioritization remains accessible to visual inspection via its blob, mean subtraction and meanDoG maps. The detected blobs, their associated records and the corresponding mean maps allow users to determine whether a high score is supported by a spatially coherent feature compatible with a plausible ligand pose. The meanDoG map generally outlines the broader volume of a localized feature, whereas the mean subtraction map localizes the strongest residual density more precisely. These representations provide related but complementary information.

We evaluated FoXSight across seven protein targets, combining enrichment analysis for six campaigns with recovery of reported M^pro^ hits and comparisons of representative maps. Where PanDDA comparison maps were available (Figs. 3–5 and Supplementary Figures S3 and S5–S9), the two approaches provided overlapping but not identical information. Agreement between the methods can increase confidence in a candidate, while differences can reveal features whose detection is sensitive to background modeling and reference selection. A particular strength of FoXSight is its robustness against structural non-isomorphism, as illustrated by the challenging Cathepsin L campaign.

Both FoXSight and PanDDA detect similar ligand-associated regions in most of the analyzed examples, while individual cases differed in the spatial extent and visual prominence of their density features. These differences are consistent with the underlying approaches for density detection.

In the BAZ2B x538 example, FoXSight recovered a weak ligand-associated feature despite competing ground-state solvent density. This ligand assignment was independently supported by anomalous density at the bromine position [10]. The selected M^pro^ examples further illustrate that the meanDoG map can provide coherent coverage of spatially demanding compounds when the corresponding conventional or event-map density appears fragmented at the displayed contour levels. These examples highlight that FoXSight facilitates identifying and interpreting weak and spatially distributed density features.

In addition, FoXSight was successfully applied as the primary hit detection tool in the screening campaign for human Cathepsin L. The analyzed datasets exhibit a continuous variation of the unit-cell parameters (Supplementary Figure S1). This non-isomorphism hampered the construction of sufficiently homogeneous ground-state ensembles for PanDDA. Also, a pre-clustering with cluster4x [11] did not fully resolve this problem because several resulting clusters retained substantial internal heterogeneity. In the analyzed cohort of 1,481 datasets, FoXSight identified 121 hits. Using cluster4x and PanDDA, 46 (38%) of these hits fell in clusters graded good, okay or medium for manual inspection.

A distinctive strength of FoXSight is its ability to operate across markedly different dataset scales. At its minimal useful configuration, the core subtraction and filtering procedure can compare a single target dataset with one suitable reference dataset. This capability enables pairwise residual-density analysis even when only a few crystals are available. However, such an analysis does not benefit from the majority-voting consensus scoring. If additional unbound or otherwise suitable reference datasets are available, independently calculated subtraction maps can be combined through voting, reducing sensitivity to the characteristics of a single reference and increasing confidence in recurrent features. The same framework can therefore support both focused experiments involving only a few datasets and screening campaigns resulting in thousands of datasets. This flexibility makes FoXSight a useful complement to multi-dataset-based approaches such as PanDDA and pairwise dataset comparison approaches as implemented in Xtrapol8. FoXSight follows a different approach and provides a scalable framework for detecting, localizing, and prioritizing residual density that can rely on a pairwise comparison, but also gain robustness as further reference datasets are incorporated.

Beyond its flexibility across dataset sizes, FoXSight’s median runtime of about 1.5 minutes per dataset on a single CPU core and straightforward parallelization support practical screening workflows. These properties also make it feasible to revisit individual datasets with alternative reference selections or parameter settings as interpretation progresses.

FoXSight’s performance could benefit from extensions in the future. Dataset-dependent parameterization could account for measurable properties such as resolution, map noise, protein size, solvent content, or the extent of structural variation. For example, the threshold used for blob growth could be adjusted according to the *σ* level of each DoG map. In addition, the geometric, map-based, and consensus-derived properties recorded for each blob provide a basis for complementing the empirical FoXSight score with a trained confidence model. Methods such as support vector machines,[23] gradient boosting[24], or neural networks[25] could integrate these features to derive a calibrated hit probability measure. However, only independent training and test campaigns can assist in evaluating generalization while controlling for protein– and dataset-level information leakage.

The FoXSight maps also pave the way to new methodological advances beyond dataset prioritization and modeling. Because the meanDoG signal often outlines the ligand-associated volume, it could provide spatial constraints for density-guided ligand placement. In a prospective modular workflow, poses generated in this manner could be optimized using methods such as JAMDA[26] and subsequently expanded using, e.g., FastGrow.[27] Such an integration would connect dataset prioritization, ligand placement, pose refinement, and fragment-to-lead design. The computational scalability of the method could likewise be improved by implementing the DoG convolutions using GPU-compatible frameworks such as CUDA[28] or OpenCL,[29] further reducing the runtime for large screening campaigns if enough GPU calculation capabilities are available.

In summary, our results establish FoXSight as a scalable and interpretable approach for hit identification in crystallographic fragment screening experiments, in particular because it can localize weak ligand-associated density. The consistent behavior using a single default parameterization across proteins of different sizes, resolutions, and structural variability indicates a useful practical operating range and highlights the robustness of the method. FoXSight’s distinctive strength is its breadth of applications: the core analysis can operate on a single target-reference pair, while additional reference datasets provide increasingly robust consensus. Moreover, the same framework remains applicable to campaigns resulting in thousands of datasets. With runtimes of a few seconds per dataset for the tested BAZ2B datasets on a current laptop, FoXSight can also be applied online during data collection, so that hits are identified while the screening experiment is still in progress. By presenting localized and transparent density-based evidence, FoXSight shifts expert effort from exhaustive searching toward focused validation and model building. Therefore, it represents both an immediately applicable addition to crystallographic screening workflows and a foundation for increasingly automated fragment-to-lead processes, encompassing ligand identification, refinement, and downstream lead compound design.

## 3 Methods

FoXSight applies multi-reference subtraction, three-dimensional DoG filtering, blob detection and consensus voting to identify candidate ligand-associated density (Fig. 1). Each target is compared with similar reference maps to highlight residual features. DoG filtering emphasizes spatial features at the scale of small-molecule fragments. Related filters are used for automatic feature detection in other life-science applications [30, 31]. Blob-level votes summarize recurring features across reference comparisons. The resulting records and maps support prioritization and visual assessment within crystallographic modeling workflows.

### 3.1 Input Data

FoXSight requires the 2*mF*_o_ *DF*_c_ electron density maps of both the target structure and one or more corresponding subtrahend structures as primary input. For Cathepsin L, these maps were calculated with phenix.mtz2map from the DIMPLE output. For the public datasets, the maps provided with the depositions were used and recalculated with Phenix where required. Optional inputs encompass a pre-refined structural model of the target (omitting density of known structures, see below), and, whenever applicable, the structure of the expected small molecule.

### 3.2 Density Map Subtraction

Initially, FoXSight performs subtraction of electron density maps, specifically, aligned 2*mF*_o_ *DF*_c_ maps, to highlight density unique to the target structure. Maps are converted to regular grid containers and the appropriate affine transformations are used to align target and subtrahend maps in real space. Trilinear interpolation ensures accurate correspondence between target and subtrahend grids, yielding:

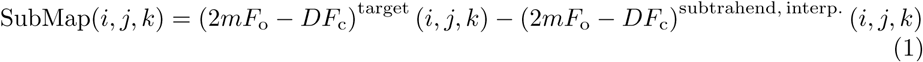

where (*i, j, k*) are the indices of the target grid, and the superscript subtrahend, interp. indicates that the subtrahend map is interpolated at the position of each target grid point. For this, each target grid point is converted to orthogonal coordinates, scaled by the ratio of the unit-cell edge lengths of the two crystals, and converted to the subtrahend grid. The subtraction map therefore lies on the target grid. No structural superposition is performed, so target and reference datasets must share the crystal form and reference frame, as obtained by refinement against a common starting model. These subtraction maps are still unsuitable for automatic detection, since peak values are prone to being noise-related.

For each candidate reference *s*, FoXSight records the subtraction residual *R*(*d, s*), the mean absolute value of the subtraction map over the grid points inside the search volume (the target model’s bounding box in all reported evaluations, see Supplementary Table S3). A target dataset is never used as its own reference. References are ranked by increasing *R*(*d, s*), and the seven with the smallest residuals are used for voting. Randomly drawn reference pools can contain hits. Their effect is limited because each reference contributes a separate subtraction map, so a reference with density at the same site reduces the support for a target feature only in that map. Their mean, *R_d_*, normalizes the dataset score. FoXSight does not rescale the maps, so *R_d_* is comparable between datasets whose maps share the same scaling, as for the sigma-scaled maps used here.

FoXSight’s default parameters, summarized in Table S3 of the Supplementary Information, are optimized for 2*mF*_o_ *DF*_c_ maps calculated in Phenix [32] with sigma scaling. Alternative map calculation methods (e.g., Gemmi [33]) may necessitate corresponding parameter adjustments.

### 3.3 Feature Enhancement and Noise Reduction via DoG Filtering

The next step in the algorithm enhances meaningful features and reduces noise. Following map subtraction, FoXSight applies a three-dimensional DoG filter to enhance blob-like features within the electron density. DoG filters are particularly sensitive to blob– or cloud-like features, a spatial motif that corresponds to atoms and chemical bonds in electron density maps. The DoG filter is a band-pass filter widely used in computer vision for feature detection [34]. It serves as an efficient approximation of the *Laplacian of Gaussian* (LoG) operator, the sum of the second-order spatial derivatives of a Gaussian. The approximation is closest for σ2 ≈ 1.6 σ1. FoXSight derives both widths from a filter radius *r* as 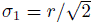 and 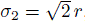, that is, *σ*_2_ = 2*σ*_1_.

Specifically, two Gaussian filters with different standard deviations, denoted by *σ*_1_ and *σ*_2_, are convolved with the subtracted map. The difference between the resulting maps is computed as:

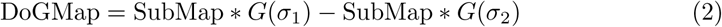

where *G*(*σ*) denotes a 3D Gaussian kernel with standard deviation *σ*, and represents convolution. The direct implementation exploits Gaussian separability by applying successive one-dimensional filters along each axis. Alternatively, symmetric convolution can be evaluated by multiplication in the discrete cosine transform (DCT) domain [35]. FoXSight’s spectral implementation uses this approach to evaluate the same sampled, truncated Gaussian kernels with reflected boundary conditions. It can reduce filtering time for larger search regions, while direct convolution can be more efficient for very small regions (e.g., a binding site). Kernel support, transform overhead and restricted-region handling are described in the Supplementary Methods.

In FoXSight, the default radius of the DoG filter is *r* = 1.75 Å (*σ*_1_ 1.24 Å, *σ*_2_ 2.47 Å). This choice is motivated by the typical spatial extent of electron density in compounds, resulting from typical inter-atomic bonds and is also used in the DoGSite method [31].

The resulting noise-reduced DoGMaps accentuate peaks in regions corresponding to candidate binding events.

### 3.4 Blob Detection via Breadth-First Search

To identify salient features in the DoG-filtered electron density maps, the highest-valued voxels are selected as seed points for further analysis. Starting from each seed point, we apply a *Breadth-First Search* (BFS) [36] over the 26-connected neighboring voxels of the map grid to iteratively group spatially connected voxels that are likely to belong to the same feature. This procedure is iterated for a user-defined number of *N_Blobs_* seed points and is set to 25 by default. We call these features blobs.

A neighboring voxel is added to the current blob if its DoG-filtered value exceeds a specified threshold *thresh_BlobSeed_*, set at 35% of the initial seed value by default. This threshold was empirically determined to provide a close approximation of the true binding compound volume while minimizing inclusion of noise or unrelated density.

### 3.5 Consensus Voting Based on Blob Similarity

Consensus voting measures whether blobs recur across subtraction maps calculated from different references. The *N*_closest_ = 7 references with the smallest mean absolute subtraction residual are used by default. Blobs from different maps are considered similar when their centroids are within 2.0 Å or their volumetric overlap exceeds 66%.

Let *B_s_* denote the blobs detected in subtraction map *s*, and *r*(*B*) the originating map of blob *B*. Its vote count is

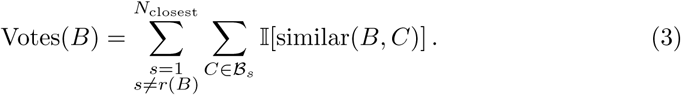

Here, I is the indicator function. The originating map contributes no votes, and votes count supporting blobs rather than distinct reference maps. Multiple blobs from one other map can therefore contribute, although counts are usually of the same order as the number of references. The blob map visualizes these spatial features and their support, while the dataset score weights each blob by its squared vote count. A high count indicates recurrence across reference comparisons, not a calibrated probability of ligand binding or a substitute for map inspection.

### 3.6 Dataset Score

To rank datasets for inspection, FoXSight first sums a contribution of every detected blob into the blob score of dataset *d*:

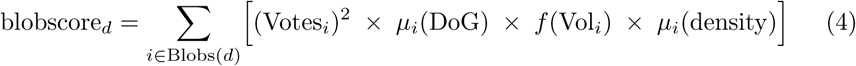

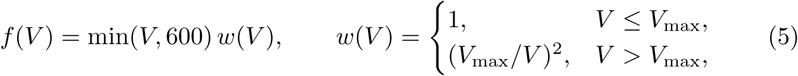

with *V*_max_ = 300 A^3^, where Votes*_i_* counts supporting blobs from other subtraction maps, *µ_i_*(DoG) is the mean DoG value, *µ_i_*(density) is the mean target-map density, and Vol*_i_* is the estimated volume of blob *i* (Supplementary Information). Up to *V*_max_, the volume term grows with *V*. It then decreases as 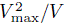 for 300 *< V ≤* 600 and as 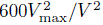 above 600 Å^3^. Thus, a 600 Å^3^ blob counts like a 150 Å^3^ blob. This weight limits the contribution of extended differences. The selected ceiling should lie above the expected ligand-associated blob volume and may need to be increased or disabled for larger compounds (Supplementary Information).

The blob score also grows with differences between a target and its references that are unrelated to ligand binding, such as lower resolution, unit-cell variation or conformational changes. Consensus voting does not remove these contributions. A difference that lies in the target appears in every subtraction map and is therefore supported by all references, like a bound ligand. A bound fragment, however, is small and localized, whereas global differences extend over the whole search volume. The subtraction residual *R_d_*, the mean residual of the seven references used for voting (see Density Map Subtraction), therefore measures this global level of difference. The dataset score divides the blob score by it, so that blobs count relative to how strongly the dataset differs from its references overall, analogous at the level of whole datasets to the per-voxel normalization of PanDDA Z-maps:

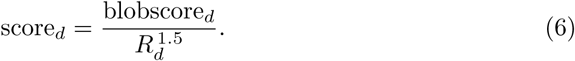

The exponent and volume ceiling are empirical choices motivated by global map agreement and the expected scale of fragment-associated features. Several blob-score factors can increase with global target–reference differences, motivating normalization stronger than division by *R_d_* alone. We use an exponent of 1.5 within a broad performance plateau between 1 and 2. FoXSight already computes *R_d_* for reference selection, so normalization requires no additional map calculations. Parameter-sensitivity and campaign-held-out checks are described in the Supplementary Information and Supplementary Figures S13 and S14. The score is a transparent prioritization heuristic. Datasets are ranked by decreasing score.

### 3.7 Output

FoXSight generates output for visualization and programmatic analysis. CCP4-formatted electron density maps marking detected blobs and mean subtraction/DoG maps can be used in molecular graphics environments. Metadata for each blob— including position, volume, and score—are exported in JSON or plain text format, enabling straightforward downstream statistical analysis and workflow integration.

### 3.8 Optional Steps

#### 3.8.1 Omit Behavior

A user-supplied pre-refined model of the target can be used to omit electron density assigned to known protein or solvent atoms, thereby focusing the analysis on novel or unmodeled density. The appropriate affine transformations are used to place all structure copies into the map. Afterward, density within a radius of 0.425 Å of any protein atom (setting 1, used in all reported evaluations) or, optionally, also of modeled small molecules and solvent (setting 2) is omitted from the subtraction map prior to DoG filtering. Mean subtraction maps are calculated without omission (Supplementary Information).

#### 3.8.2 Site-Focused Analysis

FoXSight can optionally focus the analysis on a known region of interest, such as an active site, specified by residue numbers and a chain identifier. DoG filtering is then restricted to a sphere with a radius of 13 Å around the selected site, and subsequent blob detection and consensus analysis operate only within this region. Restricting the computationally demanding filtering step to a smaller volume can substantially reduce runtime. This option was not used in the evaluations reported here, which assessed the generalization of the default parameterization across different protein targets without prior binding-site selection.

#### 3.8.3 Compound Structure

If the expected compound structure is provided as input, the blob-growing threshold may be adaptively relaxed until the detected blob matches the compound’s calculated volume. For low-occupancy hits, this can cause blobs to grow along inconsistent directions in different reference maps, so that they fail the similarity criteria. This could in theory lead to false negatives. Future work could tune the parameterization to account for this behavior.

### 3.9 Implementation and Computational Requirements

FoXSight was initially developed within the NAOMI ChemBio Suite [37]. The analyses reported here used FoXSight version 1.0.0, a standalone C++ implementation based on Gemmi [33] and RDKit [38] with a command-line interface for scripting and pipeline integration. Run metadata record the executable and source-build identifiers and analysis settings. A separate deployment is accessible through the Proteins*Plus* API (v2) [39, 40]. The benchmarks used the standalone executable.

The Gaussian filtering for the three-dimensional DoG maps is computed in the frequency domain with discrete cosine transforms. Scores agree with a direct realspace implementation up to floating-point rounding, and all rankings are identical. Runtime depends on the number of reference maps and the size and sampling of the analyzed region. Each of the 2,740 cluster analyses ran on one CPU core, with at most 75 concurrent jobs. Computation times were taken from the FoXSight logs, excluding scheduling and external file-transfer times. Peak memory was measured as maximum resident set size. All analyses ran on four identical cluster nodes, each with four Intel Xeon E5-4620 processors (eight cores each, 2.2 GHz base and 2.6 GHz maximum clock frequency). Campaign wall-clock spans were measured separately from the first analysis start to the last completion. Supplementary Methods describe the distributions and size-adjusted outlier analysis.

Median runtimes per dataset ranged across campaigns from 27 s (M^pro^) to 157 s (JMJD1B), and 95% of the analyses finished within 3.2 minutes (maximum 10.2 minutes, Supplementary Figures S10 and S11). Computations for individual targets are independent and can be parallelized across CPU cores or cluster nodes. With at most 69 and 75 concurrent single-core jobs, respectively, the 388 JMJD1B datasets were processed in 27 minutes and the 1,481 Cathepsin L datasets in 47 minutes.

For the laptop comparison, all 201 BAZ2B datasets were processed sequentially with the same arguments and reference-list order using a macOS build on an Apple M4 MacBook Pro with 16 GB RAM. The single-threaded analyses required 2 to 12 s per dataset (median 4.4 s, compared with 30 s on the cluster), with results agreeing with the cluster outputs up to floating-point rounding. Times are FoXSight’s logged computation times, as for the cluster analyses. These short runtimes support on-the-fly analysis during data collection once processed target maps and suitable references are available. They also make it practical to revisit ambiguous datasets with alternative references or settings.

Peak resident memory had a median of 1.2 GiB, with 95% of analyses below 2.1 GiB and a maximum of 5.4 GiB (Supplementary Figure S12). These per-analysis requirements are compatible with common workstations. Parallel execution requires an appropriate combined memory budget.

## Acknowledgments

This work was funded by the BMFTR (XDD Project, 13K22GUB). This research was supported in part through the Maxwell computational resources operated at Deutsches Elektronen-Synchrotron DESY, Germany. The authors acknowledge the beamline scientists at PETRA III MX beamline P11 at DESY for technical support and the group of Dǔsan Turk at the Jǒzef Stefan Institute in Slovenia for providing Cathepsin L for crystallization experiments.

## Author contributions

T.S. developed and implemented FoXSight, performed the computational analyses, and prepared the figures. M.R. and A.M. conceived the study. T.S. designed the study, selected the datasets, and developed the validation strategy. S.F. and S.G. performed the crystallographic screening and data collection and evaluated hit-finding data. S.F. processed the crystallographic data and performed model building and refinement. A.M. and M.R. supervised the research and acquired funding. T.S. wrote the initial manuscript. All authors contributed to manuscript revision and approved the final manuscript.

## Code Availability

The FoXSight source code (version 1.0.0) used for all analyses reported here will be released under the BSD 3-Clause license through the UHH-RareyLab GitHub organization (https://github.com/UHH-RareyLab) at the latest upon journal publication. Until then, a versioned package with installation instructions and example data is available from the corresponding authors upon request. FoXSight is also accessible through the Proteins*Plus* API (https://proteins.plus/api/v2) [39].

## Data Availability

Screening data for the public BAZ2B, SP100, JMJD2D, JMJD1B, and BRD1 datasets are available via Zenodo [13–17]. Scores, logs and run metadata of all 2,740 FoXSight analyses, the FoXSight maps shown in the figures, and the FoXSight inputs of all M^pro^ analyses and of the four Cathepsin L examples, including the 42 apo reference maps, are available in the accompanying data repository [41]. The complete Cathepsin L screening results will be reported in a separate publication. The Cathepsin L structures listed in Supplementary Table S1 are available from the Protein Data Bank.

## Supplementary information

The Supplementary Information provides Cathepsin L compound and structure identifiers, data collection and refinement statistics, default FoXSight parameters, additional methodological details, a runtime and memory benchmark, and further map comparisons.

