## Supplementary Information for "FoXSight: Facilitating Computational Hit Identification in Crystallographic Fragment Screening Experiments"

#### 1 Supplementary Results

Supplementary Figures S1–S5 document the structural heterogeneity of the Cathepsin L campaign and provide additional map comparisons for BAZ2B and $M^{pro}$. For BAZ2B and $M^{pro}$, five hit crystals not shown in the main text were chosen at evenly spaced ranks of the data resolution, from best to worst. The two BAZ2B hits found after cluster4x preclustering are shown separately. Conventional maps, FoXSight maps, and, where available, PanDDA maps are displayed at the contour levels described in the main text.

Representative comparisons for BRD1, JMJD1B, JMJD2D, and SP100 are provided in Supplementary Figures S6–S9. Five matched hit crystals are shown for each of the first three datasets, and both available SP100 hit crystals are included. The selection spanned three quantiles (5%, 50%, and 95%) of mean heavy-atom ligand-model occupancy and the best and worst remaining resolutions, using one site per crystal. Recorded human site choices were retained where available. Otherwise the existing site with the highest mean model occupancy was used. Following visual review, JMJD2D-x401 was replaced by x336 to provide a clearer example at the same mean model occupancy (0.22), and the worst-resolution example x568 was replaced by x463, which carries the largest ligand among the remaining crystals (14 heavy atoms). The resulting illustrative subset is not a random sample or an independent detection-performance assessment. Map-comparison PNGs for all hit crystals of these datasets, including those not selected for the Supplementary Figures, are available in the accompanying Zenodo data repository together with the selection provenance [1].

#### 1.1 Cathepsin L campaign context

**Table S1** Compound and structure identifiers for the four representative human Cathepsin L hits shown in the main manuscript. Crystal IDs are internal campaign identifiers. The abbreviated dataset labels used in the figure are given in parentheses.

| PDB ID | IUPAC name | Compound SMILES | Compound ID | Crystal ID (dataset label) |
|---|---|---|---|---|
| `pdb_000034cj` | 2-(benzimidazol-1-yl)-*N*-methylacetamide | `O=C(NC)CN1C=NC2=C1C=CC=C2` | EOS102745 | `CatL_13687_2` (13687/2) |
| `pdb_000034ck` | 5,8-dimethoxy-1,2,3,4-tetrahydroisoquinolin-4-ol | `OC1C2=C(C(OC)=CC=C2OC)CNC1` | EOS102786 | `CatL_13703_1` (13703/1) |
| `pdb_000034cl` | 3-(4-bromophenyl)-2-methylpropanamide | `O=C(N)C(C)CC1=CC=C(Br)C=C1` | EOS103020 | `CatL_12562_1` (12562/1) |
| `pdb_000034cm` | 8-hydroxy-2,6-diazaspiro[3.5]nonan-5-one | `OC(CNC1=O)CC21CNC2.Cl` | EOS103144 | `CatL_12754_1` (12754/1) |

The compound identities of the representative Cathepsin L hits are summarized in Table S1, and their data collection and refinement statistics are provided in Table S2. The campaign comprised 2,260 collected single-crystal datasets from screening of the European Fragment Screening Library. Of these, 1,481 datasets, including 42 apo datasets, passed the complete DIMPLE pipeline [2] and were potentially suitable for FoXSight analysis and model building. The crystals belonged to space group P1 and contained four protein molecules in the asymmetric unit. The ranked FoXSight cohort comprised these 1,481 datasets, with 121 confirmed-hit crystals. All 121 positive dataset identifiers match entries in the campaign hit list, each linked to a distinct compound database identifier. Thus, the 121 crystal-level positives correspond to 121 compound records, rather than repeated crystals of the same recorded compound. The unit-cell distributions in Supplementary Figure S1 summarize 1,480 of these datasets, as no cell dimensions were recorded for one dataset. The 42 apo datasets of this cohort formed the candidate reference pool. The cell distributions and model ensemble (Supplementary Figures S1 and S2) provide context for the clustering and PanDDA analyses.

***Qualitative assessment of PanDDA inspection workload.***

Following cluster4x subdivision and PanDDA analysis, clusters were assigned to five practical inspection categories: good, okay, medium, bad and really bad. This was a qualitative assessment of manual inspectability, rather than a numerical quality cutoff or a blinded sensitivity benchmark. The good, okay and medium categories were considered feasible for manual inspection and contained 46 of the 121 known hit datasets. The remaining 75 fell in the bad or really bad categories. These counts describe the location of known positives within the assessed workflow. They do not establish that PanDDA failed to detect ligand-associated density in the remaining datasets.

### 2 Supplementary Methods

Table S3 summarizes the evaluated configuration. The numerical parameters use the software defaults, whereas both bounding-box options were enabled explicitly. Unless otherwise stated, these settings were shared across analyses. Reference ensembles were selected as described in the main text.

#### 2.1 Reference-ensemble selection

For each target dataset, FoXSight ranks candidate references by their mean absolute real-space subtraction residual and uses the seven most similar maps for blob voting (Table S3). For SP100, BRD1, BAZ2B, JMJD2D and JMJD1B, the candidate pool comprised 100 randomly selected datasets. For $M^{pro}$ and Cathepsin L, all available apo datasets were considered. This keeps the evaluated numerical configuration fixed while allowing the reference pool to reflect each campaign's available data.

#### 2.2 Dataset prioritization and enrichment metrics

The empirical score ranks datasets for inspection and does not alter blob detection or map generation. Datasets were sorted by decreasing score, retaining input-table order for ties. For a top-ranked fraction $f$, recovery is the proportion of confirmed hits among the first $\lceil fN \rceil$ datasets, where $N$ is the number ranked. EF10 is the hit rate in the highest-ranked 10% divided by the overall hit rate. AP is the mean precision at ranks containing a confirmed hit. Recovery AUC was calculated by trapezoidal integration of cumulative recovery against the inspected fraction, including the origin. It is distinct from ROC AUC. AP and EF10 should be interpreted alongside hit prevalence and count, especially for SP100 with only two confirmed positives. These metrics assess prioritization against the available annotations rather than sensitivity or specificity of map interpretation. CatL annotations derive from a FoXSight-led hit-identification workflow.

The JMJD2D enrichment analysis includes all 226 completed datasets. Annotations for the original 208 datasets were retained unchanged, including 58 event-only annotations. The additional 18 datasets contribute four confirmed-hit annotations, including additional cluster4x hits [3], and 14 datasets without reported hits from the historical annotation records. No additional event-only annotations were inferred from these binary hit labels. The six-campaign enrichment cohort therefore comprises 2,715 datasets and 264 confirmed-hit datasets.

The score is a heuristic built from consensus support, density and expected feature size, with residual normalization for overall reference agreement. Its few empirical settings were chosen within broad ranges of similar performance. The initial parameter-sensitivity and campaign-held-out checks below used the original annotated cohort of 2,697 datasets and 260 confirmed hits, including 208 JMJD2D datasets. The selected score settings were retained unchanged for the expanded final evaluation. The enrichment tables and rank-movement figures report the full 2,715-dataset cohort. Blob-based scoring supported prioritization during Cathepsin L hit finding. Residual normalization and volume weighting were subsequently assessed using the available campaign annotations.

***Residual normalization.***

The reported score is the blob score, including the volume weight described below, divided by $R_d^{1.5}$. $R_d$ is the mean subtraction residual of the seven references used for dataset $d$: for each reference, the mean absolute subtraction-map value over the grid points of the search volume, as computed by FoXSight for reference selection. The residual mainly reflects global differences between a dataset and its references, such as resolution, map quality, unit-cell variation and conformational changes. A bound fragment contributes little to it, because its volume is small compared with the protein's search volume. The normalization therefore lowers datasets whose blob score is inflated by poor agreement with the references.

The exponent range was determined in the initial cohort by leave-one-campaign-out cross-validation over the six annotated campaigns. The candidates were exponents of 0, 0.5, 1, 1.5, 2, 3, 4, 6 and 8, applied to the mean residual of the seven references or, alternatively, to the residual of the best-matching reference alone. In each fold, the candidate with the highest mean normalized EF10 (EF10 divided by the best attainable EF10) on the five remaining campaigns was applied to the held-out campaign. The folds selected an exponent of 1 in five cases (three times with the mean residual, twice with the best single residual) and 2 in the remaining fold (BAZ2B held out). Across the six held-out campaigns, these selections placed 113 instead of 103 of the 260 confirmed hits in the top 10%. On all six campaigns, the objective is nearly flat between exponents 1 and 2: 0.540 at 1, 0.532 at 1.5 and 0.535 at 2, compared with 0.430 without normalization. We therefore report the mean residual with an exponent of 1.5, in the middle of this range. Exponents above 2 overcorrect: they also penalize datasets that differ from their references because of the ligand itself, for example through a ligand-induced conformational change, or that contain a hit at lower resolution (Supplementary Figure S13, BAZ2B and BRD1). Supplementary Figure S13 shows how every confirmed hit moves in the ranking for exponents between 0 and 4. Supplementary Table S4 lists the enrichment of the blob score without normalization.

***Volume weight.***

Blob volumes are estimated from the number of grid points $N$ of a blob and the mean grid spacing $s$ of the map lattice as $V = Ns/4$. This is not an exact physical volume, but for ligand-associated blobs it typically deviated by about 15–20% from the calculated compound volume. The blob score caps each blob's volume term at

$600\,\text{Å}^3$ and multiplies it by $w(V) = 1$ for $V \leq V_{\max}$ and $(V_{\max}/V)^2$ above, with $V_{\max} = 300\,\text{Å}^3$ (main-text Methods, Dataset Score). Blobs much larger than a bound fragment therefore contribute less. The weight was tested in the initial cohort after the exponent had been fixed, with the same leave-one-campaign-out cross-validation and $k = 1.5$. We compared absolute ceilings in $\text{Å}^3$, including an optional lower ceiling that down-weights small blobs. We also tested ceilings relative to compound volume and absolute ceilings whose steepness grows with $R_d$. The grid covered lower ceilings of 0 to $200\,\text{Å}^3$, upper ceilings of 100 to $1200\,\text{Å}^3$ or none, and steepness exponents of 0.5 to 8. With absolute ceilings, the held-out selections placed 131 instead of 121 confirmed hits in the top 10% (95% bootstrap interval of the change, −1 to +17), mainly in JMJD1B and BRD1. Ceilings relative to the compound volume did not transfer between campaigns (116 hits), and coupling the steepness to $R_d$ gave no further gain. The upper ceiling carries the improvement: on all six campaigns, the objective rises from 0.532 without weight to 0.588 with the best upper ceiling alone and to 0.549 with the best lower ceiling alone. A lower ceiling also moved more hits down, because some confirmed hits produce only small blobs, so none is used. Upper ceilings between 200 and $400\,\text{Å}^3$ perform almost equally, and we use $300\,\text{Å}^3$ with a steepness of 2. In the expanded final evaluation, the selected score places 134 of 264 confirmed hits in the top 10% and leaves the top-50% recovery unchanged relative to residual normalization alone (237 of 264). With the weight, exponents of 1, 1.5 and 2 place 126, 134 and 137 hits in the top 10%. We keep 1.5. Supplementary Figure S14 shows how the confirmed hits move.

$V_{\max}$ is a setting for fragment screens and must not clash with the expected ligand volume. In the reported campaigns, the median volume of the strongest blob (highest peak value) in confirmed-hit datasets lies between 112 and $270\,\text{Å}^3$, below $V_{\max}$. For larger ligands, such as lead-like compounds or peptides, the weight would also down-weight genuine binding density. $V_{\max}$ should then be set clearly above the expected ligand blob volume, for example at least at the van der Waals volume of the largest screened compound, or the weight should be disabled.

#### 2.3 Direct and spectral DoG filtering

FoXSight provides two implementations of the same discrete DoG filter. The direct implementation applies each Gaussian kernel in successive one-dimensional passes along the three axes. The spectral implementation uses the convolution–multiplication relation for symmetrically extended data [4]. A forward DCT-II is applied to the orthogonal working grid after sampling and structural omission. The transformed values are multiplied by the difference between the responses of the two separable Gaussian kernels, and an inverse DCT-III returns the filtered map. The responses are calculated from the same sampled, truncated kernel weights as in the direct implementation, rather than from the analytic transform of an infinite continuous Gaussian. This preserves equivalence with direct filtering up to floating-point rounding. The kernel weights are not additionally normalized. Half-sample reflection at the working-grid boundaries preserves the direct real-space implementation's boundary behavior without introducing periodic wraparound.

For restricted analyses, the transformed region contains the requested output region together with a surrounding halo covering the larger Gaussian kernel's finite support. Artificial crop boundaries therefore lie beyond the support of retained output values. Where the crop reaches an original working-grid boundary, the original reflection rule is retained. Transform dimensions can be expanded within the existing grid to lengths with small prime factors for computational efficiency. This includes additional existing samples without introducing padding or resampling. Values outside the requested output region are set to zero before downstream analysis.

Spectral filtering can substantially reduce computation time for larger filtering regions and broader Gaussian kernels. Restricting the transform to the selected region and its halo avoids transforming the complete working grid. However, direct convolution can be more efficient for very small regions, where transform overhead and the required halo outweigh the computational savings. The crossover depends on grid dimensions, kernel support and hardware, and the speed advantage need not increase monotonically with map size. Even small regions on large maps can benefit because the direct implementation retains full-grid intermediate arrays. Sampling, other full-grid operations and downstream analysis remain, so filtering speedups do not translate directly into equivalent reductions in total runtime or memory. The implementation does not automatically select a method based on search-region size. For typical kernel sizes, we recommend the spectral filter.

Numerical comparisons used synthetic fields, including constant inputs and boundary impulses, and real crystallographic maps with full-grid and restricted filtering. The tests checked preservation of the finite-kernel response and boundary behavior, together with agreement of map values and downstream blob records. Observed differences between direct and spectral evaluation were limited to floating-point rounding in the tested cases. All runtime measurements reported in this manuscript used the spectral implementation.

#### 2.4 Search-volume restriction

FoXSight can optionally restrict the analysis volume using the `--boundingbox` command-line option. When this option is used with a target PDB or mmCIF model supplied through `--pdb`, a bounding box is constructed around the heavy atoms of the protein (solvent and ligands excluded) and expanded by 2.0 Å. Map subtraction and reference scoring are then restricted to this box. The additional `--DoGonlyBB` option also restricts the DoG output, and therefore blob detection, to the box. Both options were enabled for all evaluations reported in this manuscript.

Restricting the search volume can reduce distracting or redundant features outside the protein-centered region. It does not itself denoise the retained map values. However, symmetry-related protein copies outside the box may contain genuine ligand-associated features, potentially with clearer density than the copy within the box. Very large ligands or binding sites remote from the supplied protein model may also extend beyond this region. If relevant features approach the box boundary or appear better resolved around another symmetry mate, the analysis can be repeated without bounding-box restriction. For the screening datasets considered here, the protein-centered search volume encompassed the relevant ligand-binding regions.

#### 2.5 Site-focused analysis

Residue numbers and a chain identifier can be supplied to focus FoXSight on a known site. With this option, DoG filtering is computed only within a sphere of radius 13 Å around the selected site. The subsequent blob detection and consensus steps are consequently confined to the same region. Reducing the volume processed by the DoG filter can substantially reduce runtime, but ligand-associated features outside the sphere are not assessed by the focused analysis.

Site-focused analysis was disabled for all evaluations reported in this manuscript to assess the generalization of the same default parameters across protein targets without prior binding-site selection. The reported runtime measurements therefore do not include the additional speed benefit of site-focused filtering.

#### 2.6 Runtime benchmark

Runtimes were recorded for all 2,740 analyses across seven campaigns using the configuration in Table S3, including both bounding-box options. The timing cohort includes all 2,715 datasets from the enrichment analysis and 25 $M^{pro}$ datasets used only for reported-hit recovery and map comparisons. Each analysis was an independent single-threaded job on one CPU core, with at most 75 concurrent jobs. Computation time was taken from the FoXSight log, excluding scheduling and external file transfers. Peak memory was maximum resident set size. All jobs ran on four identical cluster nodes, each with four Intel Xeon E5-4620 processors (eight cores each, 2.2 GHz base and 2.6 GHz maximum clock frequency) and 252 GiB memory. The nodes were shared with other users' jobs. Relative to runtime expected from map size, median runtime per node ranged from 0.95 to 1.02 times the expected value. The benchmark used the build with frequency-domain DoG filtering (Methods).

Median runtimes ranged from 27 s for $M^{pro}$ to 157 s for JMJD1B, with a median of 89 s over all analyses (Supplementary Figure S10). Summed single-threaded computation time was 72.3 hours. This is a sum of logged elapsed times, not a separate measurement of CPU utilization. Observed campaign wall-clock spans ranged from approximately 2 minutes for the 25 $M^{pro}$ datasets (at most 11 concurrent jobs) to 47 minutes for the 1,481 CatL datasets (at most 75 concurrent jobs). These spans include the effect of scheduling and parallel execution. Theil–Sen fits of computation time against output-map voxel count, used as a map-size proxy, gave log–log slopes from 0.74 (JMJD2D) to 0.85 (BAZ2B and $M^{pro}$), indicating sublinear scaling over the observed ranges (Supplementary Figure S11). Fixed costs such as reference-map loading can contribute to slopes below one.

To identify unusually slow analyses, residuals of these fits were converted to robust $z$-scores using the median absolute deviation. Runs were flagged when $z > 3.5$ and their computation time exceeded their campaign's median. The latter condition prevents small, fast maps from being flagged solely because fixed costs are not represented in the log–log fit. Nineteen of the 2,740 analyses (0.7%) were flagged, 1.1 to 1.7 times slower than expected for their map size. Nine were CatL analyses that ran on one node within 28 minutes, and three were JMJD1B analyses that ran on another node within 17 minutes, consistent with temporary load on the shared nodes. The other seven were

SP100 analyses on three nodes. SP100 runtimes follow map size particularly closely, so slowdowns of 8 to 44% already exceeded the threshold. Excluding the flagged analyses changes no campaign median by more than 3 s, so the medians and 95th percentiles above describe the expected runtime. Two analyses were flagged by the marginal criterion, which applies the same robust threshold within each campaign without size adjustment: JMJD1B x1292 (613 s), which has the largest map of all analyses (52 million voxels), and BAZ2B x513 (69 s). Both are consistent with their map size.

Peak memory per analysis ranged from 0.21 GiB to 5.4 GiB, with a median of 1.2 GiB, and 95% of the analyses required less than 2.1 GiB (Supplementary Figure S12). Dataset medians ranged from 0.35 GiB for BAZ2B to 1.4 GiB for JMJD1B and Cathepsin L. Like runtime, memory grows with map size: the largest values occurred for JMJD1B and Cathepsin L, and only three of the 2,740 analyses exceeded 4 GiB.

#### 2.7 Voting scheme

Candidate blobs satisfying the plausibility criteria (maximum DoG response $\geq 0.011$, volume $\geq 10\,\text{Å}^3$, and at most 10,000 grid points) are compared across subtraction maps calculated with different references. Two blobs support one another when their centroids are within 2 Å or their overlap exceeds 66%. The originating map contributes no votes. Votes count individual supporting blobs, so several similar blobs from one other map can contribute. They are not a count of distinct references or a calibrated binding probability. The dataset score uses the squared blob-level count defined in the main-text Methods.

#### 2.8 Omit behavior

When a pre-refined PDB or mmCIF model of the target is supplied, three omit settings can be selected. Setting **0** omits no density. Setting **1** omits density within a 0.425 Å radius of each protein atom and its symmetry mates. Setting **2** additionally omits density around pre-refined small molecules, including metals and solvent molecules, within the same radius.

Omission is applied after map subtraction and before DoG filtering. The selected radius is approximately one quarter of the van der Waals radius of carbon, allowing FoXSight to suppress density immediately adjacent to the pre-refined model while retaining sensitivity to larger structural changes.

The omit setting can also affect sensitivity. Masking density around pre-refined ligands can reduce model-associated background features, but it can suppress meaningful ligand-associated signal when the existing model overlaps the true binding event. For example, in BAZ2B-x538, a pre-refined solvent molecule overlaps the ligand-associated density. Masking density around such modeled solvent atoms can obscure the DoG signal needed for blob detection, consistent with the corresponding example in the PanDDA supplementary information [5]. Before using omit mode 2, the model supplied for omission should therefore be inspected for solvent molecules or pre-existing ligands that overlap potential binding sites. For ambiguous sites, results can be compared with omit mode 1 or with an appropriately prepared omission model. This concerns the model used for density masking and does not imply that legitimate solvent molecules should be removed from the final refined structure.

##### 2.9 Number of Blobs

The number of blobs retained per subtraction map, $N_{\mathrm{Blobs}}$, is an important meta-parameter because it determines which features are available for overlap analysis and voting. A suitable value depends on ligand occupancy and on the number of solvent molecules with variable positions in the crystal. The value $N_{\mathrm{Blobs}} = 25$ was used throughout the presented experiments and provided robust performance across the analyzed datasets. Lower values may suffice for datasets such as BAZ2B. Conversely, in the SP100-x596 dataset, $N_{\mathrm{Blobs}} = 10$ did not retain a high-ranking ligand-associated blob, whereas the consensus blob obtained with $N_{\mathrm{Blobs}} = 25$ closely followed the modeled ligand.

##### 2.10 Contour levels and solvent–ligand separation

For all map comparisons, the molecular-graphics carve was kept at 2.5 Å. Unless stated otherwise, conventional $2mF_{\mathrm{o}} - DF_{\mathrm{c}}$ maps were displayed at $1.0\sigma$ and $mF_{\mathrm{o}} - DF_{\mathrm{c}}$ maps at $\pm 3.0\sigma$. PanDDA event maps were displayed at $2.0\sigma$ and Z-maps at raw Z-scores of $\pm 4$, the level used in the original PanDDA publication [5]. FoXSight meanDoG and mean subtraction maps were displayed at $3.0\sigma$ and $2.0\sigma$, respectively. These levels provide a consistent visual comparison but are not detection thresholds. Displaying the meanDoG map at $3.0\sigma$ can produce large connected features when strong ligand-associated signal is spatially connected to solvent density. Adjusting the contour level can help distinguish such components during manual interpretation. This can be useful during model building when the mean subtraction map alone does not delineate the full contour of the binding molecule.

In the representative public-dataset figures, columns show conventional $2mF_{\mathrm{o}} - DF_{\mathrm{c}}$ density (blue) and positive/negative difference density (green/red), FoXSight meanDoG (blue) and mean subtraction (green), and PanDDA event density (blue), with original Z-map values (green/red) where available. Original Z-values are displayed at $\pm 4$ without re-normalization. Event-density contours at two stored-grid standard deviations are a display convention, not reconstructed Z-scores or significance levels equivalent to the other map types. For JMJD1B, the PanDDA column contains only event density reconstructed from deposited event-map coefficients. Original Z-maps were unavailable. Each row uses the same camera across methods, gray ligand carbons and protein, and 2.5 Å carving. Crystal IDs, PDB accessions and resolutions are indicated. Full site identifiers remain in the selection metadata.

#### 3 Best Practices and Troubleshooting

1. **Inspect the omission model.** Before using omit mode 2, check for modeled solvent or pre-existing ligands that overlap potential binding sites. Masking density around these atoms can suppress weak ligand-associated signal before DoG

filtering. For ambiguous sites, compare with omit mode 1 or an appropriately prepared omission model (Section 2.8).

2. **Check the bounding-box coverage.** Bounding-box restriction can exclude distracting or redundant features, but also genuine signal around symmetry-related protein copies outside the box. If a candidate lies near the boundary or another symmetry mate has clearer density, consider repeating the analysis without this restriction (Section 2.4).
3. **Revisit plausible events that lack a detected blob.** Absence of a retained blob does not establish absence of binding. Omission settings, search-volume restriction, reference selection, and the number of retained blobs can affect whether weak features are detected (Section 2.9).
4. **Inspect complementary maps together.** Blob maps localize consensus features, meanDoG maps show their broader extent, and mean subtraction maps localize residual density more directly. No omission is applied to mean subtraction maps, making them useful for assessing structural changes. A high dataset score alone does not validate a ligand assignment.
5. **Inspect blob maps in increments of 10.** Blob-map values are calculated as $v_{\text{blob}} = \text{votes} \cdot 10 + v_{\text{voter id}} + \max(\text{DoG})$.
6. **Keep map preparation consistent.** The default parameters were selected for sigma-scaled Phenix $2mF_{\text{o}} - DF_{\text{c}}$ maps. Alternative map-generation or scaling procedures may require parameter adjustments (Table S3).
7. **Facilitate comparisons between runs.** Retaining the reference-dataset identifiers, software version, and settings used can be helpful when comparing analyses, particularly when exploring alternative references or parameter choices.

### 4 Supplementary Discussion

#### 4.1 Combinatorial approaches

Reference selection provides a natural point of interaction with existing crystallographic methods. For example, Cluster4x [3] could construct more specialized reference ensembles by grouping datasets according to crystallographic and structural similarity. Reference datasets could then be selected separately for individual targets or groups of closely related targets. Such a strategy could reduce systematic subtraction artifacts and increase the relevance of the reference ensemble. However, highly specialized ensembles may contain too few suitable reference datasets or retain internal heterogeneity, as observed for parts of the Cathepsin L campaign. The balance between structural similarity, ensemble size, and reference diversity therefore merits systematic investigation.

A second strategy is to combine FoXSight with PanDDA-derived evidence. PanDDA Z-maps and event maps are not equivalent to FoXSight meanDoG or mean subtraction maps, so numerical values should not be combined without normalization and validation. Instead, spatial overlap with PanDDA-supported regions, or the presence of a corresponding PanDDA candidate, could provide additional evidence for FoXSight voting or scoring. Joint analysis could prioritize regions supported by both approaches while retaining each map for interpretation.

**Table S2** Data collection and refinement statistics for the four representative human Cathepsin L hits. Values in parentheses refer to the highest-resolution shell. The X-ray wavelength was 1.033 22 Å. The criteria used to determine the maximum resolution were $R_{\mathrm{pim}} \leq 0.6$, $I/\sigma I \geq 2$, and $\mathrm{CC}_{1/2} \geq 0.3$.

| Crystal ID | `CatL_13687_2` | `CatL_13703_1` | `CatL_12562_1` | `CatL_12754_1` |
|---|---|---|---|---|
| Compound | EOS102745 | EOS102786 | EOS103020 | EOS103144 |
| PDB code | `pdb_000034cj` | `pdb_000034ck` | `pdb_000034cl` | `pdb_000034cm` |
| **Data collection** | | | | |
| Space group | P1 | P1 | P1 | P1 |
| $a$ (Å) | 57.1303 | 56.9988 | 57.3958 | 57.0797 |
| $b$ (Å) | 62.6811 | 62.5081 | 62.6693 | 62.6612 |
| $c$ (Å) | 67.7871 | 67.3258 | 67.9440 | 67.6046 |
| $\alpha$ (°) | 105.3040 | 105.3769 | 105.5241 | 105.3801 |
| $\beta$ (°) | 93.5813 | 93.7880 | 93.4874 | 93.7973 |
| $\gamma$ (°) | 115.5933 | 115.5228 | 116.1137 | 115.5197 |
| No. unique reflections | 69753 | 57654 | 34350 | 81333 |
| Resolution (Å) | 53.50–1.79 (1.82–1.79) | 53.40–1.91 (1.95–1.91) | 64.16–2.27 (2.31–2.27) | 53.51–1.70 (1.72–1.70) |
| $R_{\mathrm{merge}}$ | 0.099 (0.746) | 0.133 (0.826) | 0.179 (0.672) | 0.140 (0.682) |
| $I/\sigma I$ | 12.3 (2.3) | 13.2 (2.2) | 8.9 (2.5) | 9.7 (2.4) |
| Completeness (%) | 92.0 (88.0) | 94.5 (88.7) | 92.3 (84.6) | 92.2 (86.1) |
| Redundancy | 6.1 (4.6) | 10.4 (6.3) | 5.2 (4.8) | 7.4 (4.2) |
| $\mathrm{CC}_{1/2}$ | 0.998 (0.703) | 0.997 (0.767) | 0.989 (0.787) | 0.994 (0.712) |
| **Refinement** | | | | |
| Resolution (Å) | 53.50–1.79 (1.82–1.79) | 53.32–1.91 (1.94–1.91) | 53.15–2.27 (2.33–2.27) | 53.45–1.70 (1.72–1.70) |
| No. unique reflections | 69739 | 57638 | 34350 | 81333 |
| $R_{\mathrm{work}}$ | 0.1597 (0.2694) | 0.1690 (0.2348) | 0.1633 (0.2072) | 0.1680 (0.2583) |
| $R_{\mathrm{free}}$ | 0.2041 (0.3426) | 0.2215 (0.3192) | 0.2211 (0.2674) | 0.2034 (0.2853) |
| No. atoms | | | | |
| All | 7718 | 7527 | 7490 | 7734 |
| Ligand | 28 (14 per mol.) | 15 | 13 | 22 (11 per mol.) |
| Water | 560 | 491 | 470 | 601 |
| $B$-factors | | | | |
| Protein | 23.08 | 27.15 | 28.33 | 21.83 |
| Water | 27.67 | 29.36 | 29.95 | 26.77 |
| Other | 42.51 | 45.51 | 43.69 | 40.57 |
| R.m.s. deviations | | | | |
| Bond lengths (Å) | 0.007 | 0.008 | 0.004 | 0.006 |
| Bond angles (°) | 0.80 | 0.82 | 0.55 | 0.77 |
| Ramachandran outlier | 0 | 0 | 0 | 0 |

**Table S3** FoXSight configuration used in the reported evaluations, comprising the numerical defaults and explicitly enabled search-volume restrictions. The primary map filename records the blob-growth threshold, number of initial blobs and number of reference maps. For example, `..._T-350_BN25_NOFSUB7.ccp4`. Optional settings are distinguished from the evaluated configuration.

| Stage | Parameter and value | Definition or filename encoding |
|---|---|---|
| Map preparation | Sigma-scaled Phenix $2mF_\mathrm{o} - DF_\mathrm{c}$ maps | Map type for which the defaults were selected. Alternative map-calculation procedures may require adjustment. |
| DoG filtering | `--dog` = 1.75 Å | DoG filter radius $r$. The two Gaussian widths are $\sigma_1 = r/\sqrt{2}$ and $\sigma_2 = \sqrt{2}\,r$. |
| DoG filtering | `--mindelta` = 0.4 Å | Minimum spacing of the temporary orthogonal grid on which the DoG filter is evaluated. Filtered values are projected back to the map grid for blob detection. |
| Blob growing | `--threshold` = −0.35 | Negative values specify a seed-relative threshold. −0.35 retains voxels above 35% of the initial seed value. Encoded as `T-350`. |
| Blob growing | `--blobn` = 25 | Number of highest-valued DoG voxels used as blob seeds per subtraction map. Encoded as `BN25`. |
| Reference voting | `--nofSubs` = 7 | Maximum number of globally most similar reference maps used for voting. Encoded as `NOFSUB7`. |
| Candidate filtering | Maximum DoG response $\geq 0.011$ | Minimum peak response required before blob comparison. |
| Candidate filtering | Volume $\geq$ 10 Å$^3$. $\leq$ 10,000 grid points | Basic size and extent criteria for candidate blobs to filter out dust and large, typically noise-induced blobs, which can cause runtime outliers. |
| Consensus voting | Centroid distance $\leq$ 2.0 Å or overlap $>$ 66% | Criteria by which blobs from different subtraction maps support one another. |
| Structural omission | `--omitBehaviour` = 1, if a target model is supplied | Omits density within 0.425 Å of protein atoms and their symmetry mates before DoG filtering. Setting 0 disables omission. Setting 2 also omits pre-refined small molecules. |
| Search volume | `--boundingbox`: enabled in all reported evaluations (disabled by default) | Restricts map subtraction and reference scoring to the bounding box of the protein's heavy atoms, extended by 2.0 Å. Requires a target model (`--pdb`). |
| Search volume | `--DoGonlyBB`: enabled in all reported evaluations (disabled by default) | Additionally restricts the DoG output, and therefore blob detection, to the same box. Implies `--boundingbox`. |
| Site-focused analysis | Disabled in the reported evaluations. Radius 13 Å when enabled | Residue numbers and a chain identifier specify a local site. DoG filtering and subsequent detection and consensus analysis are restricted to the corresponding sphere. |

**Table S4** Confirmed-hit enrichment of the blob score without residual normalization (exponent 0) and without volume weight, for comparison with main-text Table 1. Metrics and ranking as in the main text.

| Target | $N$ | Full hits | Top 10% ($n/N$) | Top 50% ($n/N$) | EF10 | Recovery AUC | AP |
|---|---|---|---|---|---|---|---|
| BAZ2B | 201 | 11 | 9/11 | 11/11 | 7.83 | 0.907 | 0.453 |
| BRD1 | 302 | 43 | 21/43 | 41/43 | 4.76 | 0.832 | 0.509 |
| JMJD2D | 226 | 29 | 9/29 | 22/29 | 3.05 | 0.694 | 0.301 |
| SP100 | 117 | 2 | 0/2 | 2/2 | 0.00 | 0.697 | 0.049 |
| JMJD1B | 388 | 58 | 12/58 | 43/58 | 2.06 | 0.640 | 0.237 |
| CatL | 1,481 | 121 | 54/121 | 118/121 | 4.44 | 0.855 | 0.342 |
| Pooled | 2,715 | 264 | 105/264 | 237/264 | – | – | – |

### 5 Supplementary Figures

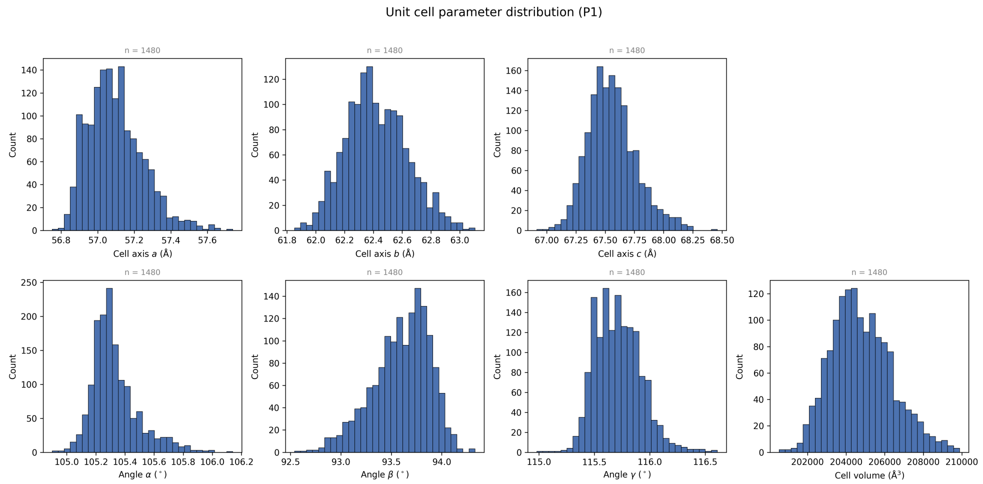


**Fig. S1** Distribution of unit-cell parameters across 1,480 of the 1,481 Cathepsin L datasets in space group P1. The broad distributions, particularly in cell volume, illustrate the non-isomorphism encountered in this screening campaign.

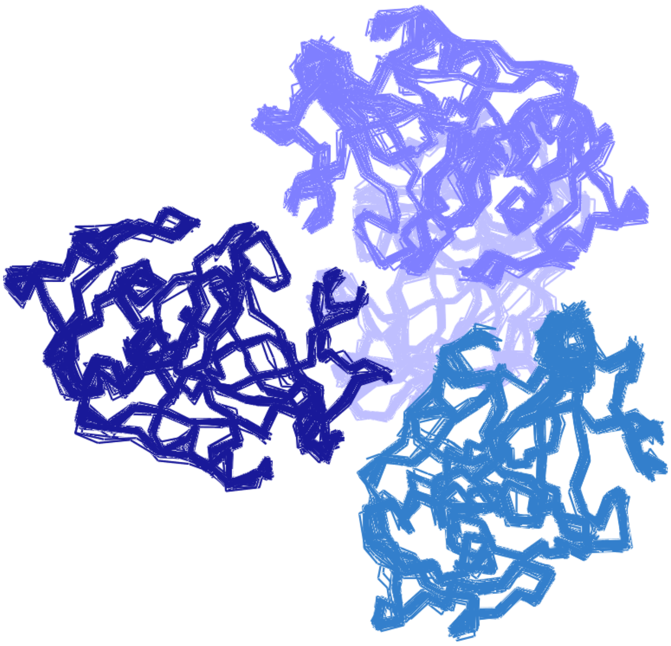

**Fig. S2** Superposition of Cathepsin L protein models from the screening campaign. The structural ensemble illustrates the conformational and packing variability present across the datasets. Colors are used to distinguish individual models.

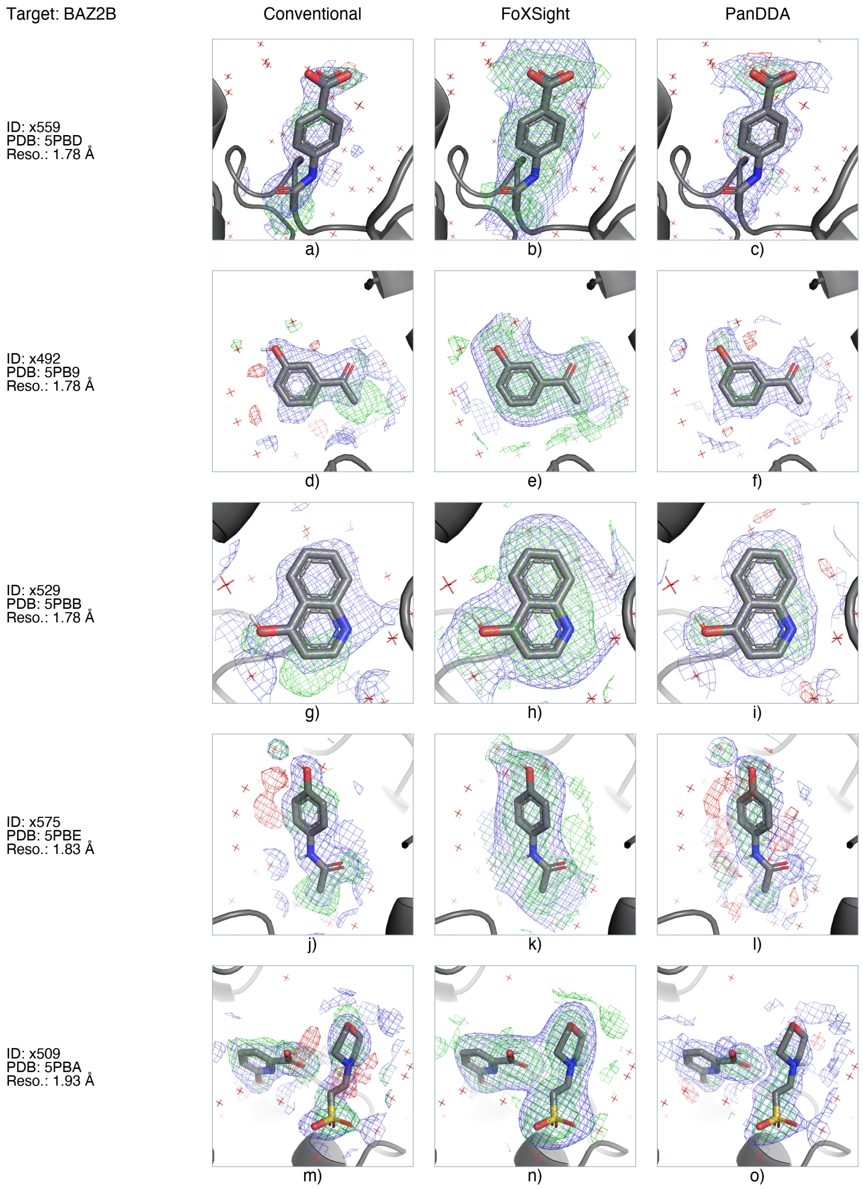


**Fig. S3** Additional BAZ2B map comparisons for five hit crystals not shown in the main text, chosen at evenly spaced resolution ranks. Columns show conventional $2mF_\mathrm{o} - DF_\mathrm{c}$ density (blue) with positive and negative $mF_\mathrm{o} - DF_\mathrm{c}$ difference density (green and red), FoXSight meanDoG (blue) and mean subtraction (green) maps, and PanDDA event (blue) and Z-map (green and red) density. Dataset identifiers, PDB accessions and resolutions are indicated in each row. Contour levels and the 2.5 Å carve radius are specified in the Supplementary Methods. Map comparisons for all hit crystals are available in the accompanying Zenodo data repository [1].

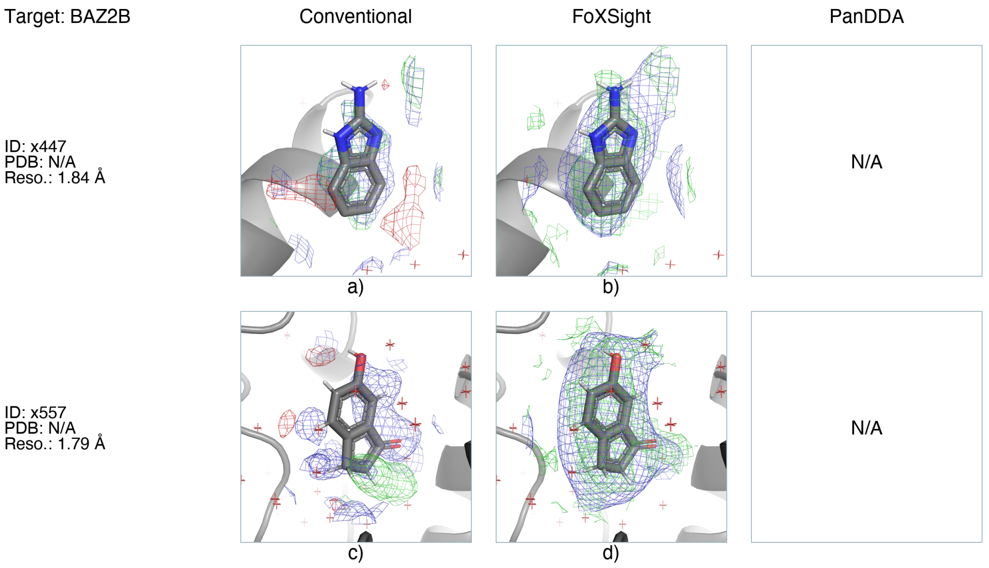


**Fig. S4** Additional BAZ2B hits identified using cluster4x preclustering before PanDDA analysis [3]. Columns show conventional $2mF_\mathrm{o} - DF_\mathrm{c}$ (blue) and $mF_\mathrm{o} - DF_\mathrm{c}$ (green/red) maps and FoXSight meanDoG (blue) and mean subtraction (green) maps. FoXSight reveals ligand-associated density for these hits, including x557, for which the conventional maps are difficult to interpret. These hits were absent from the original PanDDA hit set. PanDDA maps from the cluster4x-assisted analysis were not available among the comparison inputs used here and are therefore not shown. Their absence does not indicate that PanDDA failed after preclustering.

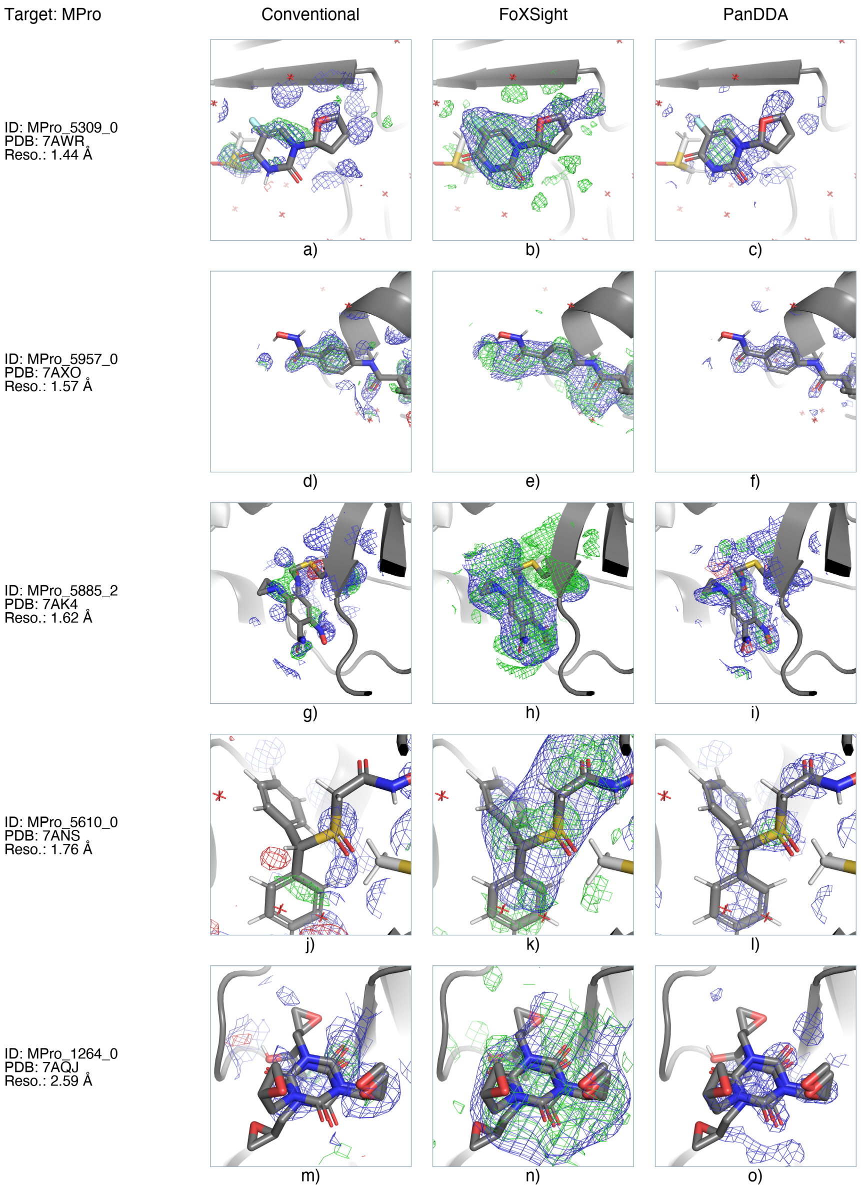


**Fig. S5** Additional $M^{pro}$ map comparisons for five hit crystals not shown in the main text, chosen at evenly spaced resolution ranks. Columns show conventional $2mF_o - DF_c$ density (blue) with positive and negative $mF_o - DF_c$ difference density (green and red), FoXSight meanDoG (blue) and mean subtraction (green) maps, and PanDDA event (blue) and Z-map (green and red) density. Dataset identifiers, PDB accessions and resolutions are indicated in each row. Available PanDDA maps are shown for individual binding events. These comparisons do not represent a uniform campaign-wide PanDDA analysis. Map comparisons for all hit crystals are available in the accompanying Zenodo data repository [1].

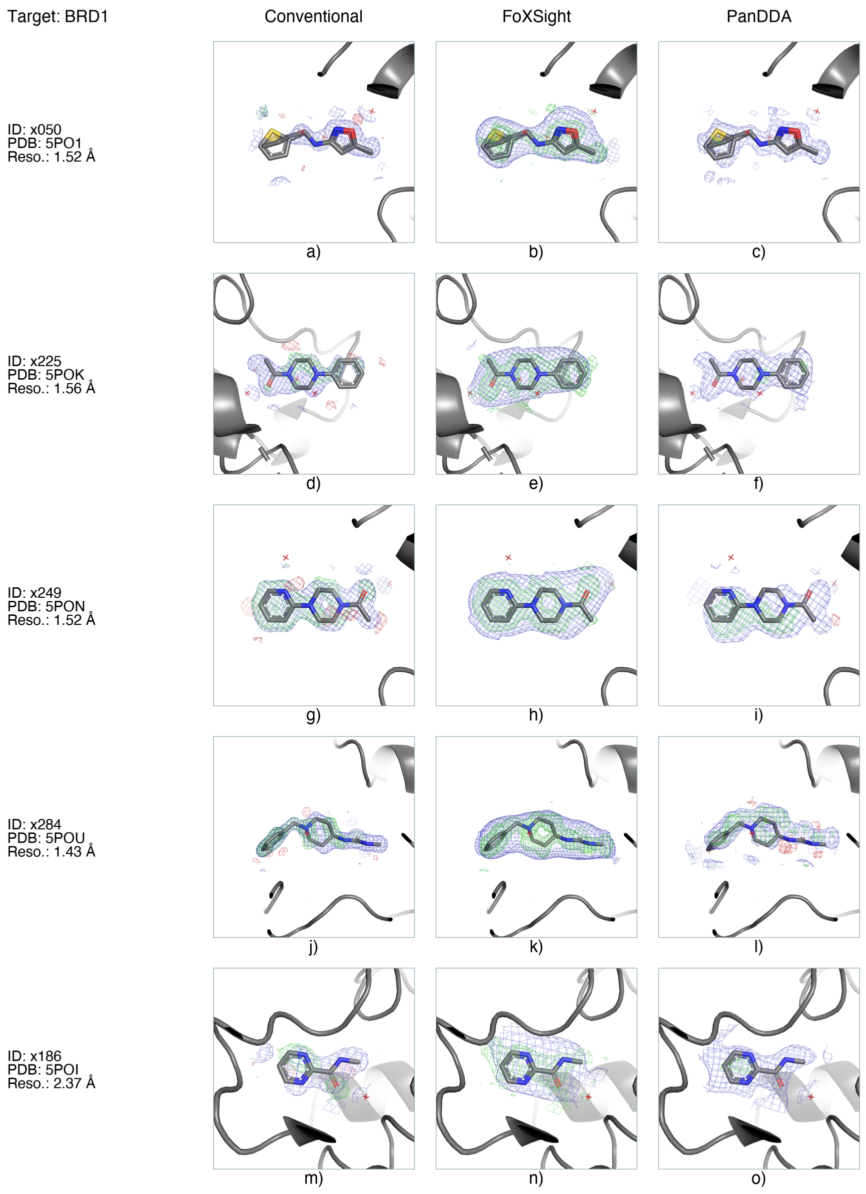


**Fig. S6** Representative BRD1 map comparisons for five selected hit crystals. Columns show conventional, FoXSight and PanDDA maps at the fixed contour settings specified in the Supplementary Methods. Each row shares one camera across methods. Crystal IDs, PDB accessions and resolutions are shown. The selection procedure is described in the Supplementary Results.

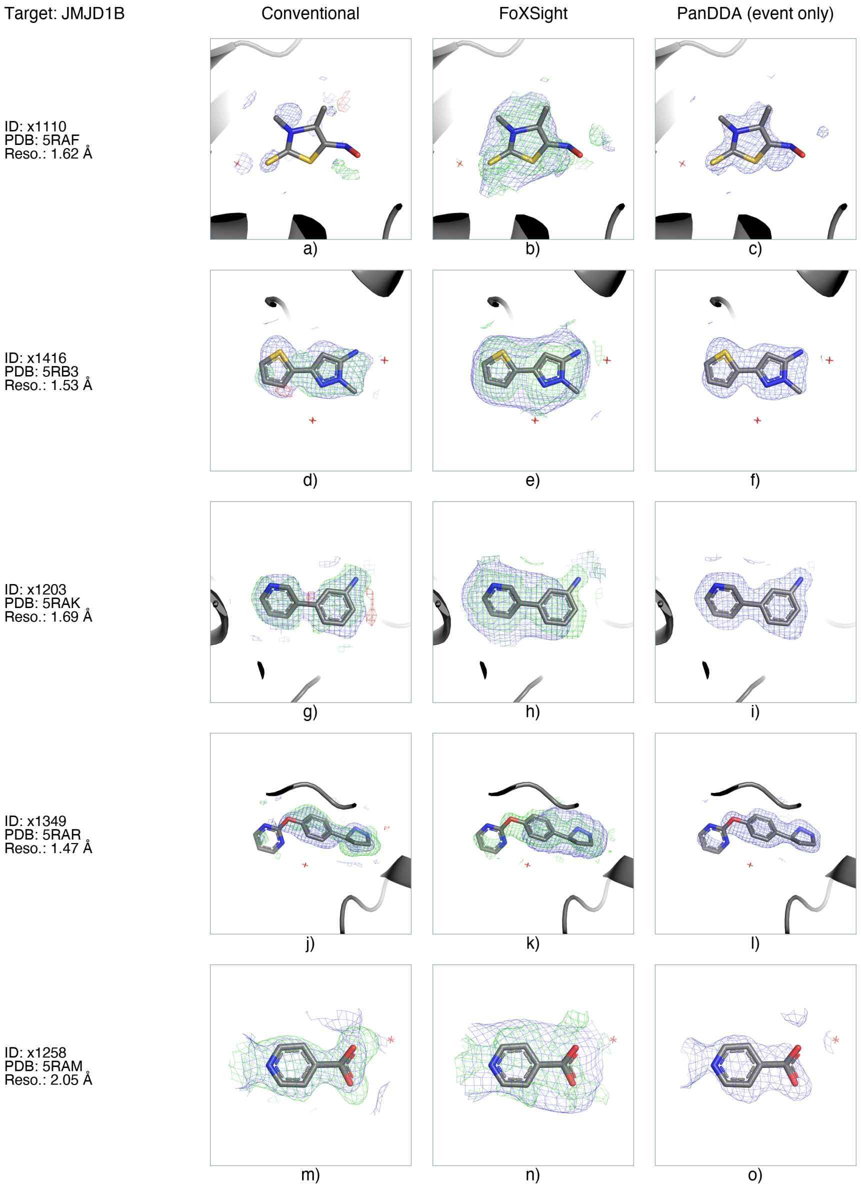


**Fig. S7** Representative JMJD1B map comparisons for five selected hit crystals. Map colors and contour settings are as specified in the Supplementary Methods. The PanDDA column contains deposited event density only. Original Z-maps were unavailable. Event-density contours are not reconstructed Z-scores.

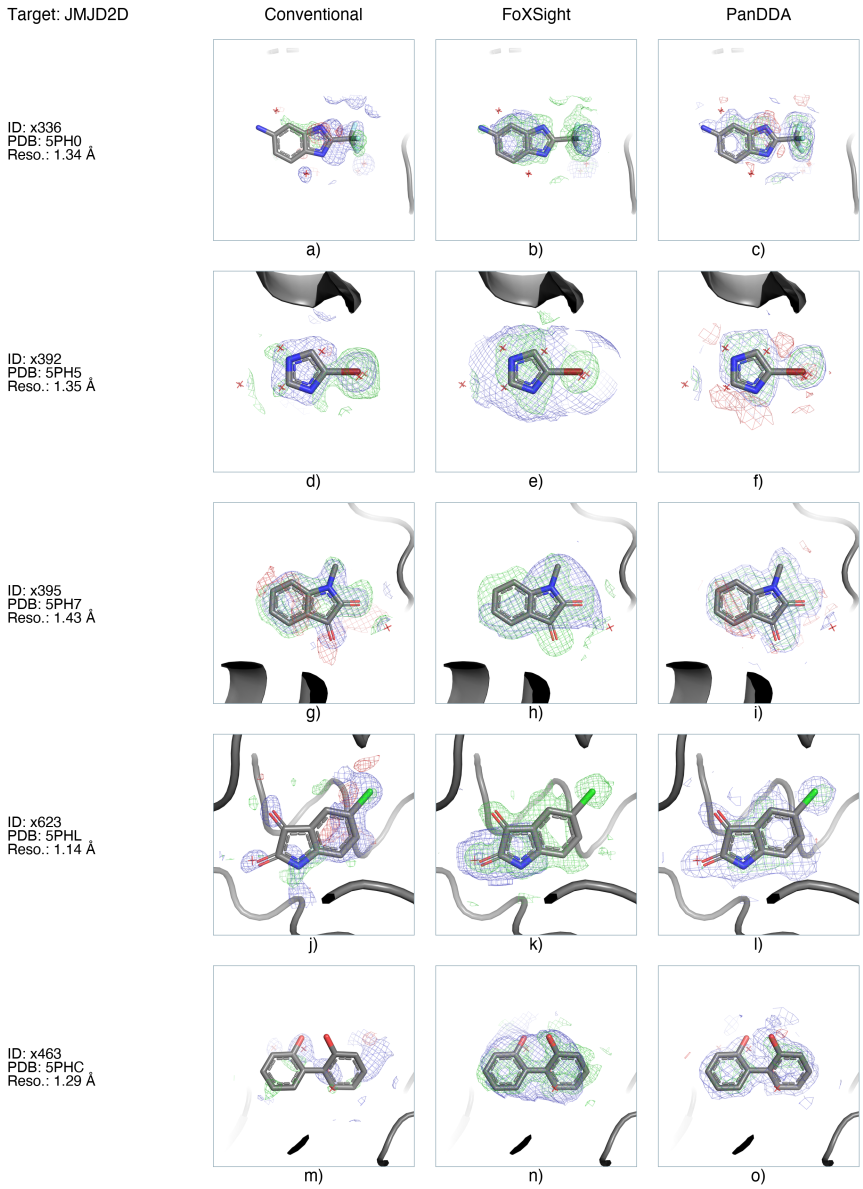


**Fig. S8** Representative JMJD2D map comparisons for five selected hit crystals. Map colors and contour settings are as specified in the Supplementary Methods. The visually selected examples x336 and x463 replace x401 and x568 of the initial automatic selection. The complete comparison archive retains all four crystals.

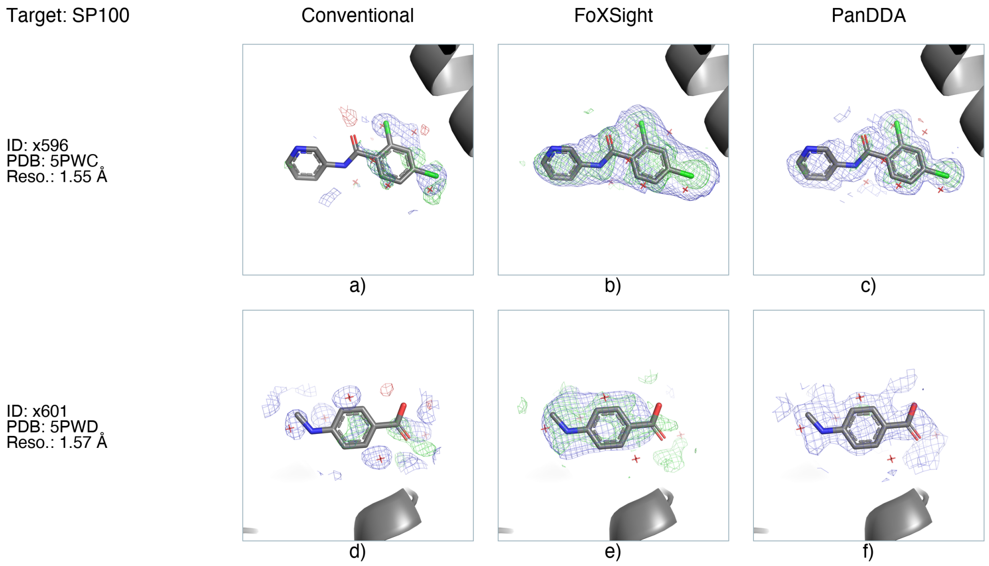


**Fig. S9** Map comparisons for both available matched SP100 hit crystals. Columns show conventional, FoXSight and PanDDA maps at the fixed contour settings specified in the Supplementary Methods. One site per crystal is illustrated. Alternative site rows are retained in the complete comparison archive.

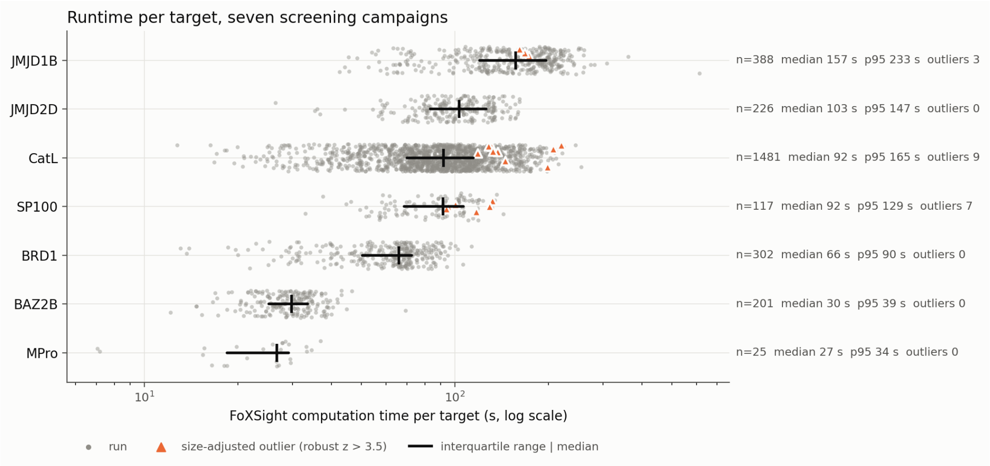


**Fig. S10** FoXSight computation time per target dataset for all 2,740 analyses of the seven screening campaigns (log scale). Each gray point is one single-threaded analysis on one CPU core. Black bars show the interquartile range and the median. The number of analyses, the median, the 95th percentile and the number of size-adjusted outliers are given for each campaign. Orange triangles mark analyses that were slow for their map size (robust $z > 3.5$, see Supplementary Figure S11). Details are given in the Supplementary Methods (Runtime benchmark).

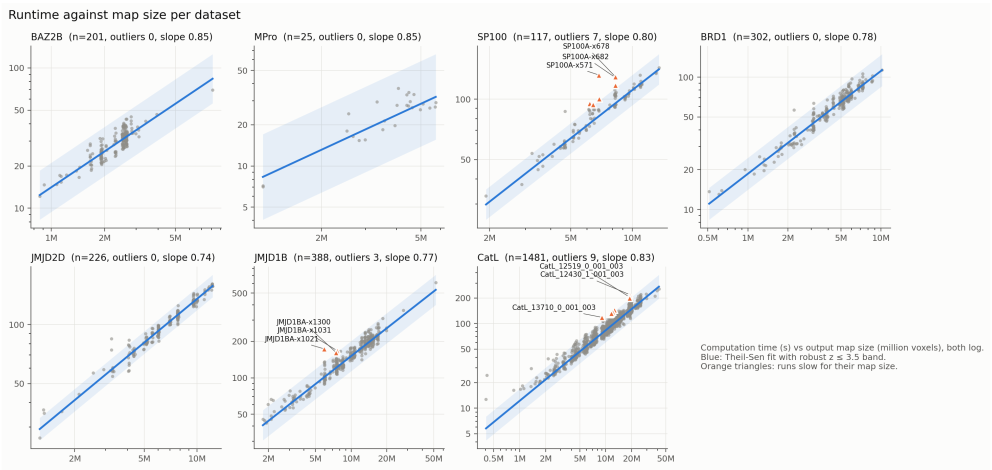


**Fig. S11** FoXSight computation time against output-map size for each screening campaign (both axes logarithmic, map size in million voxels). Blue lines show Theil–Sen fits in log–log space, and the shaded bands mark robust $z \leq 3.5$ around the fit. The fitted slope is given in each panel title. A slope of 1 corresponds to runtime proportional to map size. Orange triangles mark the 19 analyses that were slow for their map size. Details are given in the Supplementary Methods (Runtime benchmark).

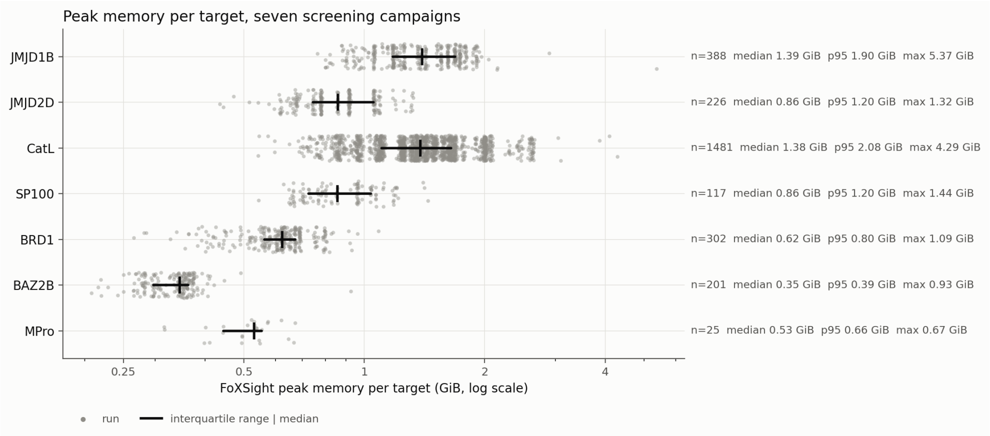


**Fig. S12** Peak memory (maximum resident set size) per target dataset for all 2,740 FoXSight analyses of the seven screening campaigns (log scale). Each gray point is one single-threaded analysis. Black bars show the interquartile range and the median. The number of analyses, the median, the 95th percentile and the maximum are given for each campaign. Campaigns are ordered as in Supplementary Figure S10.

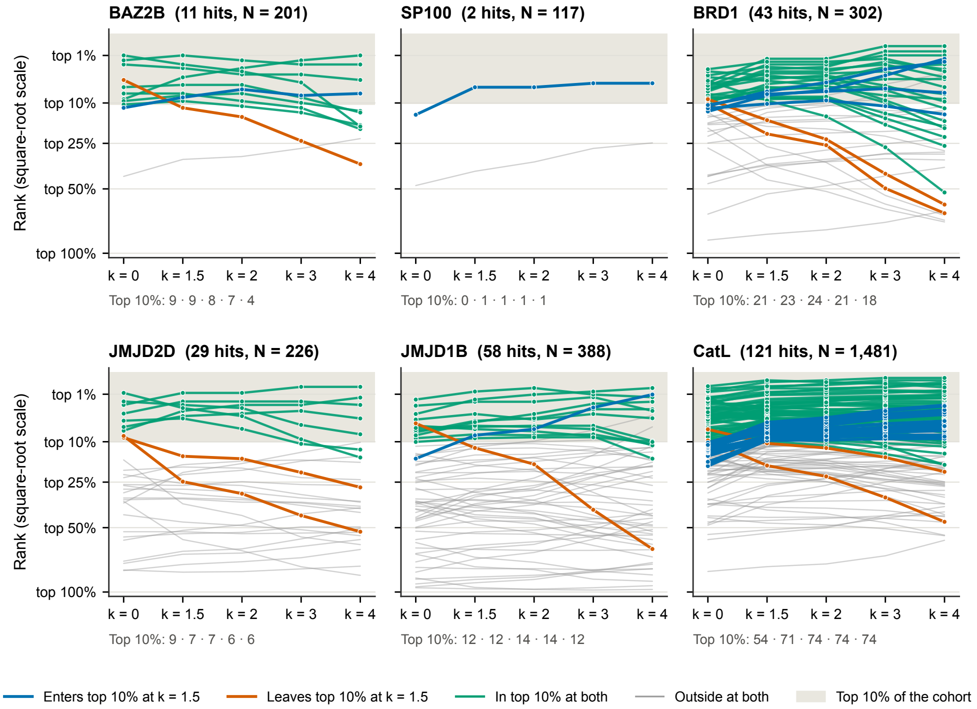


**Fig. S13** Rank of every confirmed hit under the residual-normalized score blobscore$_d/R_d^k$, without the volume weight, for exponents $k$ between 0 (blob score alone) and 4. Each line is one confirmed hit in the expanded evaluation cohort, including all 226 JMJD2D datasets. Ranks are given as the top fraction of the campaign on a square-root axis, and the shaded band marks the top 10% ($\lceil 0.1N \rceil$ datasets). Colors compare $k = 0$ with the reported $k = 1.5$: hits that enter the top 10% (blue), leave it (orange), stay in it (green) or stay outside (gray). The line under each panel gives the number of confirmed hits in the top 10% for each $k$. In BAZ2B and BRD1, exponents above 2 move hits out of the top 10% again.

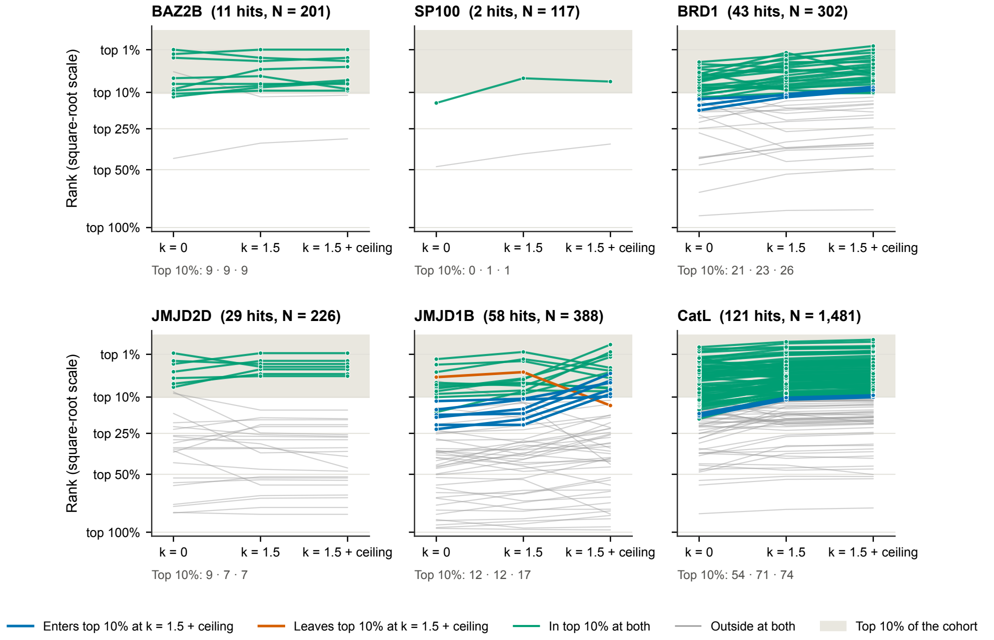


**Fig. S14** Rank of every confirmed hit under the blob score alone ($k = 0$), the residual-normalized score without volume weight ($k = 1.5$) and the reported score with the volume weight ($k = 1.5 +$ ceiling, $V_{\max} = 300\,\text{Å}^3$). The expanded evaluation cohort, axes and shading are as in Supplementary Figure S13. Colors compare $k = 1.5$ without and with the volume weight: hits that enter the top 10% (blue), leave it (orange), stay in it (green) or stay outside (gray). The line under each panel gives the number of confirmed hits in the top 10% for each column.
